# Molecular Logic of Cellular Diversification in the Mammalian Cerebral Cortex

**DOI:** 10.1101/2020.07.02.185439

**Authors:** Daniela J. Di Bella, Ehsan Habibi, Sun-Ming Yang, Robert R. Stickels, Juliana Brown, Payman Yadollahpour, Fei Chen, Evan Z. Macosko, Aviv Regev, Paola Arlotta

**Author notes:** These authors contributed equally (Daniela J. Di Bella, Ehsan Habibi). These authors contributed equally. Correspondence should be addressed to and.

## Abstract

The neocortex has an unparalleled diversity of cell types, which are generated during development through a series of temporally orchestrated events that are under tight evolutionary constraint and are critical for proper cortical assembly and function. However, the molecular logic that governs the establishment and organization of cortical cell types remains elusive, largely due to the large number of cell classes undergoing dynamic cell-state transitions over extended developmental timelines. Here, we have generated a comprehensive single-cell RNA-seq and single-cell ATAC-seq atlas of the developing mouse neocortex, sampled every day throughout embryonic corticogenesis. We computationally reconstruct developmental trajectories across the diversity of cortical cell classes, and infer the gene regulatory programs that accompany their lineage bifurcation decisions and their differentiation trajectories. Finally, we demonstrate how this developmental map pinpoints the origin of lineage-specific developmental abnormalities linked to aberrant corticogenesis in mutant animals. The data provides the first global picture of the regulatory mechanisms governing cellular diversification in the neocortex.

## INTRODUCTION

The mammalian cerebral cortex is composed of a diversity of cell types, which has expanded over the course of evolution to sustain complex cerebral functions, such as sophisticated motor behavior, sensory processing, and cognition. All these cell types are produced in a constrained temporal sequence, which in the mouse begins with progenitor expansion at around E11.5, followed by the sequential generation of excitatory neuron types (E12.5-E16.5) and the production of glia (E18.5-P1)(Greig, Woodworth et al. 2013, Lodato and Arlotta 2015). During this same time period, cell types produced outside the dorsal telencephalon, such as cortical interneurons, microglia, subpopulations of oligodendroglia, and blood vessel constituents, migrate into the cortex and populate its parenchyma (Reemst, Noctor et al. 2016, Karakatsani, Shah et al. 2019).

Substantial effort has been dedicated to understanding how cortical cell types emerge during development, and, further, how interactions between diverse cells build higher-order cortical anatomy and connectivity. Decades of prior work has elucidated the roles of specific genes during the development of some cell types, and has also clarified lineage relationships between progenitors and selected progeny (Greig, Woodworth et al. 2013, Lodato and Arlotta 2015). However, large gaps in knowledge remain. What are the global regulatory mechanisms governing the differentiation of the multiple cell types present in the neocortex? When is neuronal subtype identity established? How are lineage bifurcation decisions controlled? How might malfunctions in this process relate to disease? What is needed to address these questions is a comprehensive view of the development of all cortical cells, across all developmental times, with a molecularly defined, global logic of cellular diversification in the neocortex.

Here, we have built a first comprehensive single-cell transcriptional and epigenetic molecular atlas of the developing cerebral cortex within the prospective somatosensory field, capturing development of all cortical cell types on each day of mouse corticogenesis. We identify global longitudinal molecular changes (“molecular recipes”) that accompany fate acquisition and maturation of individual cell types, and regulatory mechanisms that define lineage bifurcations as corticogenesis unfolds. We report on the comprehensive molecular changes associated with dynamic cell transitions during multi-lineage specification, and demonstrate how this map enables identification of the mechanisms behind developmental abnormalities in cellular diversification during corticogenesis in mutant animals.

## RESULTS

### Comprehensive single-cell transcriptional atlas of the developing cerebral cortex

To generate a single cell reference atlas of the developing cerebral cortex, we profiled by single cell RNA-seq (scRNA-seq) cells that were sampled from the full thickness of the developing somatosensory cortex from mouse embryos over the entire period of corticogenesis: E11.5 (transition from amplifying to neurogenic progenitors); E12.5 and E13.5 (birthdate of layer 6 and 5 excitatory neurons); E14.5, E15.5 and E16.5 (birthdate of layer 2 to 4 excitatory neurons); and E18.5 and P1 (transition from neurogenesis to gliogenesis) (Figure 1A). Overall, we collected 78,503 scRNA-seq profiles, with 4,000 to 10,000 cells per time point (Supplementary information Figure 1A,B). We clustered the cells by a graph-based algorithm (Figure 1 B, Methods), and annotated the clusters *post hoc* by expression of known marker genes (Figure 1B and Supplementary information Figure 1 C,E and 2A). We assigned an annotation to 85 to 98% of the cells per time point, corresponding to all known cell types of the developing cerebral cortex, including rare populations, such as microglia, that represent as few as 0.3% of all profiled cells.

The scRNA-seq profiles displayed a gradual progression from progenitors to different types of cortical projection neurons and glial cells (Figure 1B,D). The earliest stages of corticogenesis were largely composed of two main classes of progenitor cells, apical (AP: *Sox2, Pax6* and *Hes5)* and intermediate (IP: *Eomes, Neurog2* and *Btg2)* progenitors (Supplementary information Figure 1D; AP+IP=80% at E11.5, 69% at E12.5, 66% at E13.5). The proportion of progenitors decreased with time, gradually replaced (from E14.5) by the different types of cortical projection neurons (PN: *Neurod2, ß-tubulin III,* and *Neurod6).* These included corticofugal projection neurons (CFuPN, divided into corticothalamic projection neurons, CThPN, and subcerebral projection neurons, SCPN), as well as different callosal projection neuron (CPN) populations (Figure 1B,D and Supplementary information Figure 2A). The two progenitor populations and their neuronal progeny formed a continuous progression rather than discrete clusters, consistent with previous studies of cell-type transitions during corticogenesis being accompanied by modest and gradual shifts in gene expression(Yuzwa, Borrett et al. 2017). IPs were present at all ages, supporting the model that both early- and late-born neurons can be generated through indirect neurogenesis (Vasistha, Garcia-Moreno et al. 2015, Mihalas, Elsen et al. 2016).

Consistent with the timing of interneuron migration to the pallium, starting at E13.5 we detected ventrally-generated cortical inhibitory interneurons, marked by the expression of *Dlx1, Dlx2,* and *Dlx5,* and glutamate decarboxylases 1 and 2 (Figure 1B,D, green, Supplementary information Figure 1C and 2B,C). Medial ganglionic eminence (MGE)-derived interneurons, expressing *Sst, Npy, Lhx6, Sox6,* and *Nxph1/2,* emerged at E13.5, while interneurons expressing *Pax6, Sp8, Cxcl14,* and *Htr3a* (caudal ganglionic eminence (CGE)-derived) first appeared at E15.5 (Supplementary information Figure 2B,C). This is in line with the sequential birthdate of MGE- and CGE-derived interneurons(Batista-Brito and Fishell 2009, Mayer, Hafemeister et al. 2018). Starting at E18.5, we detected another population of *Htr3a*-positive cortical interneurons, putatively derived from the pallial-subpallial boundary and marked by *Meis2, Etv1,* and *Pbx1* (Frazer, Prados et al. 2017), reflecting the successive invasion of the cortex by interneurons of different origins (Supplementary information Figure 2B,C).

Oligodendrocyte precursor cells (OPC: *Olig1, Olig2, Pdgfra)* and astrocytes *(Apoe, Aidhi11, Slc1a3*) were first observed at E18.5, coinciding with the end of the neurogenic period (Figure1 B,D). In addition to cells of neural origin, we identified non-neural cell types including microglia *(Aif1, Tmem119),* red blood cells (RBC: *Hb-s, Car2, Hemgn),* endothelial cells *(Cldn5* and *Mcam),* pericytes *(Cspg4, Pdgfrb),* and vascular and leptomeningeal cells (VLMC: *Col1a1, Vtn, Lgals1).*

Next, we merged the cells from all developmental time points (Figure 1C, Methods), highlighting a main differentiation continuum extending from apical progenitors towards projection neurons and glial cells. Cell types that do not originate from dorsal progenitors were excluded from the main trajectories of dorsally-derived lineages, including ventrally-derived interneurons, microglia, and cells of the vasculature and meninges. Cajal-Retzius cells, which originate from progenitors in the early medial cortex (Bielle, Griveau et al. 2005), were first detected at E11.5, and emerged from *Wnt8b-*positive medial progenitors present at that age (Figure 1C). Oligodendrocyte precursor cells (OPC) grouped closely with astrocytes and did not express the dorsal marker *Emx1*, supporting the finding that the first cortical oligodendrocytes derive from ventral progenitors (Kessaris, Fogarty et al. 2006)

Finally, we related cell identities to their spatial organization in the cortex using Slide-seq (Rodriques, Stickels et al. 2019, Stickels, Murray et al. 2020) at E15.5, a time when the bulk of neuronal cell types have already been born (Figure 1E). We scored the Slide-seq data with cell type expression signatures, comprised of the top 40 differentially expressed genes for each of the main cell types (Supplementary information Figure 3A,B). The spatial expression patterns of each cell type signature were consistent with the known position of the corresponding cell types (Figure 1E and Supplementary information Figure 3C), validating the molecular signatures from scRNA-seq.

Our data collectively provide a thorough, single-cell, molecular atlas of the developing neocortex that expands our knowledge of cortical development along with the dynamic molecular changes that accompany its cellular diversification.

### Single-cell molecular reconstruction of differentiation trajectories of cortical cell types

The molecular mechanisms driving cellular diversification in the cortex remain poorly understood. The roles of a modest number of individual genes have been uncovered (Greig, Woodworth et al. 2013, Lodato and Arlotta 2015), but a global molecular picture of all cell types through time is missing.

To study the differentiation continuum, we computationally inferred comprehensive differentiation trajectories for all cortical cell classes from the scRNA-seq atlas (Figure 2A,B). First, we performed diffusion map analysis and inferred a pseudotime of the cells along a differentiation axis (Figure 2A). The inferred pseudotime largely recapitulated the cell-type gradients identified in the low dimensionality manifold, suggesting that the main source of variation in the data emerges from the cells’ stage of development and the age of collection. Next, we applied our trajectory-inference algorithm URD (Farrell, Wang et al. 2018) to generate a tree based on the transcriptional similarity of pseudotime-ordered cells. URD randomly walks along the pseudotemporally ordered cells from a set of user-defined tips (differentiated cell types) towards a pre-defined root (common progenitors). We defined the root as progenitors present at E11.5, and tips as terminal subclasses of neurons and glia at P1. Because we only sampled progenitors of the dorsal ventricular zone, we excluded differentiated cells that originate in the ventral telencephalon or outside the brain (as their root is not available in the data). The final branched trajectory tree was built using 350,000 random walks from each tip (Figure 2B).

The inferred tree was well aligned with the cells’ differentiation status, real age (collection time), known cell subtype relationships, and expected expression of known markers (Figure 2B, Supplementary information Figure 4A-D). Undifferentiated APs preceded IPs, and neuronal and glial progeny appeared last (Figure 2B, Supplementary information Figure 4A,B). The tree correctly organized cell classes, for example grouping corticofugal neurons (SCPN and CThPN) into neighboring branches (Figure 2B). Expression of known cell type-specific markers followed the appropriate lineage trajectories (Supplementary information Figure 4D).

The inferred tree also uncovered previously unappreciated expression patterns of genes traditionally considered lineage-restricted, which we validated by Slide-seq. For example, *Rorb,* a marker of layer 4 stellate neurons, and *Fezf2*, a selector gene for SCPNs, were also expressed in the astroglia lineage, consistent with recent reports that *Fezf2* is expressed in astrocytes (Bayraktar, Bartels et al. 2020) (Supplementary information Figure 5B). *Pcp4,* a marker for CFuPN (Arlotta, Molyneaux et al. 2005), was expressed in migratory neurons of both the CFuPN and CPN lineages, which we confirmed by presence of *Pcp4* signal in the intermediate zone using Slide-seq (Supplementary information Figure 5A).

Notably, the tree showed progenitors diverging as early as E13.5 into branches terminating in glial and neuronal identities (Supplementary information Figure 4A). Based on the differentially-expressed genes between APs from these two branches, this first branchpoint seems to distinguish (i) APs enriched for *Btg2* (Florio and Huttner 2014), *Neurog2,* and *Hes6* (Jhas, Ciura et al. 2006), reflecting a potentially primed neurogenic state; and (ii) “naïve” APs expressing higher levels of radial glia markers such as *Fabp7, Dbi,* and *Slc1a3,* as well as proliferation-associated genes (Supplementary information Figure 4H). In a 3D force-directed layout embedding of the tree, the first branchpoint forms a broad gradient of cells between these two contiguous AP states (2D visualization shown in Supplementary information Figure 4H). This supports a model where APs gradually transition to an astrocytic identity (Malatesta and Gotz 2013) but may still be recruited to become neurons by neurogenic cues.

### Post-mitotic diversification of projection neurons from multipotent progenitors

While recent studies have proposed that the transcriptional profile of apical progenitors changes through time as they generate the different PN subtypes (Telley, Agirman et al. 2019), it remains debated whether fate-restricted progenitors exist (Franco, Gil-Sanz et al. 2012, Guo, Eckler et al. 2013, Gao, Postiglione et al. 2014, Llorca, Ciceri et al. 2019). We set to address this using our lineage tree.

In the tree, neuronal populations emerged via a shared molecular trajectory from one common progenitor branch. To further examine this, we first sub-clustered and embedded only the AP populations (Supplementary information Figure 4F). Although they expressed key genes associated with CFuPN (*e.g*. *Fezf2*, *Tle4*, *Bcl11b*) and CPN (*e.g. Cux1*, *Pou3f3*, *Tle1*) identities, as previously reported(Zahr, Yang et al. 2018), neither AP sub-clusters, nor their embedding pattern followed the expression of these markers, which were broadly expressed across the APs (Supplementary information Figure 4G). This argues against the presence of strictly pre-committed progenitors. Notably, while APs express the markers broadly (including co-expression of key markers like Fezf2 and Pou3f3 in the same cells), the patterns were not identical, with subtle opposing gradients in the strength of expression of *e.g. Fezf2 vs*. *Pou3f3* that may further suggest skewing rather than strict pre-commitment. Finally, combining progenitors across all time points revealed a continuum of cells ordered according to the age of collection (Supplementary information Figure 4F), rather than distinct subtypes. Thus, our data largely support a model in which apical progenitors continuously and gradually evolve while generating the distinct PN types (Yuzwa, Borrett et al. 2017, Telley, Agirman et al. 2019), with flexible exploration of the transcriptional program space of PNs.

Our analysis suggested that the diversification and identity acquisition of neuronal subpopulations occur post-mitotically. In both the low dimensionality embedding and the tree, distinct neuronal progenies progressively separated at the level of post-mitotic neurons, rather than progenitors (Figure 1 B,C, 2B and Supplementary information Figure 4C). These newborn neurons acquired a broad class identity before further differentiation into terminal subtypes (Figure 2B). For example, at E13.5, post-mitotic neurons expressed genes broadly marking all CFuPN (Supplementary information Figure 4D-E), while specific markers for CThPN or SCPN appeared around E14.5. Of note, while most PN populations became distinguishable within a few days of their birth, layer 4 *Rorb* positive stellate neurons became distinct from CPN populations only later, at perinatal stages (Figure 1D), probably reflecting a role for perinatally-incoming thalamic input in imprinting sensory modality identity (De Leon Reyes, Mederos et al. 2019).

### Distinct transcriptional programs define fate-specification trajectories of individual cortical types

To define gene programs that act across the full differentiation trajectory from progenitors to terminal cell types, we used our reconstructed tree to map dynamic transcriptional changes in gene cascades, over the development of each lineage (Figure 2C and Supplementary information Figure 6–8, Methods). At the early stages of their development, all terminal neuronal types expressed shared genes (Figure 2C), including the downregulation of cell-cycle and DNA damage-related genes *(Gadd45g),* the transient expression of neurogenic transcription factors *(Neurog1, Neurog2)* and migration-associated genes *(Sstr2, Neurod1),* and the upregulation of pan-neuronal genes *(Neurod2, Dcx, Tubb3).* Conversely, cell typespecific programs appear post-mitotically as neurons migrate away from the germinal zones, and included both well-known and novel lineage-specific marker genes. For example, for SCPN, these comprised known marker genes, such as *Bcl11b, Sox5, Thy1, Tcerg1l, Crym,* and *Ldb2,* as well as genes not previously associated with SCPN specification, such as *Pex5l* and *Fam19a1,* among others (Figure 2C and Supplementary information Figure 6). The cascade for layers 2&3 CPN showed restricted expression of CPN markers *Cux1, Satb2, Pxna4* and *Cux2,* as well as novel markers *Ptprk* and *Fam19a2* (Figure 2C and Supplementary information Figure 7). We validated the age- and cell-type specificity of newly-identified marker genes by comparison to *in situ* hybridization data from the Allen Developing Mouse Brain Atlas (Thompson, Ng et al. 2014), or by comparison to temporal transcriptional profiles of PN subpopulations from the DeCoN database (Molyneaux, Goff et al. 2015) (Supplementary information Figure 5E). Surprisingly, the neuropeptides *Npy* and *Cck,* typically found in cortical interneurons, were also detected in PN lineages, in layers 5 and 6 CPN and at lower levels in CFuPN, also validated by Slide-seq (Supplementary information Figure 5C,D). This likely represents transient neuropeptide expression, as only layers 5 and 6 CPN retain *Npy* in adult mice (Allen Cell Types Database). Similar to neurons, astrocytic lineages sharply downregulated genes associated with DNA replication and proliferation, such as *Gmnn,* while upregulating astrocytic genes like *Slc1a3, Gfap,* and *Sparcl1* (Supplementary information Figure 8). Ependymal cells also emerged directly from AP, and showed a distinct upregulation of cilia-related genes, such as *Foxj1*, *Mcidas,* and *Wdr34*, as well as genes not previously associated with this cell type, such as *Rsph4* (Supplementary information Figure 8).

Complex biological processes, such as cellular diversification, can be more robustly described by the joint activity of gene programs (“modules”) than by expression of individual genes(Dixit, Parnas et al. 2016). We therefore identified gene modules across all cell populations at each time point by non-negative matrix factorization (NMF, Farrell, Wang et al. 2018), and then annotated modules of co-varying genes with biological functions according to their top-ranked genes. We chained modules from consecutive time points based on overlapping genes (Farrell, Wang et al. 2018), to define “genetic programs” representing different aspects of corticogenesis (Figure 2D and Supplementary information Figure 5F, Methods).

While some programs were associated with broad developmental events such as radial glia identity, neurogenesis, and neuronal migration, others were specific for individual lineages (Figure 2D). Neuronal lineage-specific programs became distinguishable at E14.5, consistent with the timing of neuronal divergence in the branching tree (Figure 2B,D), supporting the model of an initial shared developmental trajectory that diverges post-mitotically. Modules associated with radial glia were inter-connected with modules associated with astrogenesis, suggesting that these cell types share highly similar transcriptional programs (Figure 2D) modulated over time.

Collectively, the data identify gene expression cascades and modules that accompany identity acquisition across multiple cell lineages, and represent a valuable resource to inform future study of neocortical development.

### Molecular codes of branchpoint transitions during corticogenesis

As transcriptional programs diverge over time, lineages are separated into increasingly finely-partitioned cell populations. The combinations of genes that control these decisions in the cortex are largely unknown. The branching tree emerging from our data offers an opportunity to identify genes accompanying lineage bifurcations that may represent factors controlling or implementing these events.

To this end, we examined gene expression over a narrow pseudotime window spanning cell lineage divergence, comparing cells before and after each branchpoint (Figure 2E, dotted circles). We first identified genes that distinguished the cells in the daughter branches from each other or distinguished cells from daughter *vs*. parent branches, as well as genes whose expression is correlated with pseudotemporal progression during this time window (Methods). Next, we trained a gradient boosting decision tree algorithm to classify and assign an importance score (mean squared error impurity with Friedman’s improvement score) to each gene. We ranked genes based on their importance score, and selected the top 10 transcription factors (TFs) and DNA-binding proteins for each daughter branch as the most likely drivers of downstream changes in cell fate (Figure 2E).

The genes associated with the branchpoints (Figure 2E) included several TFs that were previously known to functionally govern cell identity acquisition, such as *Bcl11b* and *Fezf2* in the CFuPN branch, and *Satb2* and *Pou3f2* in the CPN branch (Figure 2F). The analysis also revealed novel candidate regulators of fate specification, such as *Hivep2* for CFuPN, *Chgb2* for CThPN and putative near-projecting neurons, and *Msx3* and *Satb1* for layer 4 stellate neurons. Together, the data provides a first compendium of genes associated with the main points of lineage divergence in the neocortex, and presents candidates for future functional studies.

### Cell type-specific epigenomic landscape during neocortical development is congruent with the expression landscape

To further dissect the regulatory processes driving transcriptional changes associated with lineage segregation, we next profiled single-cell chromatin accessibility using the assay for transposase-accessible chromatin using sequencing (scATAC-seq) at three key time points: E13.5, E15.5 and E18.5 (Figure 3A). We inferred gene activity scores from the scATAC-seq data (calculated from gene body and promoter accessibility) and-annotated cell types by the activity of marker genes, including AP, IP, PN, inhibitory neurons, and glia (Figure 3A, Supplementary information Figure 9A. We next co-embedded the scATAC-seq and scRNA-seq data in a shared UMAP space, relying on gene activity and expression scores, respectively, for each gene (Methods). Data from both modalities were highly interleaved (Figure 3A, insets), indicating that chromatin accessibility captured the full cell-type spectra identified by expression, and vice versa.

Next, we used the scATAC-seq gene activity scores to build a developmental trajectory tree (Figure 3B), excluding cell types that do not arise from dorsal progenitor cells, using URD. We defined tips as the most mature cell types collected at E18.5 (Figure 3A), and calculated 200,000 random walks per tip along the pseudotime-ordered cells. In the resulting tree, cells progressed in pseudotime according to both the age of collection and their differentiation state (Figure 3B), indicating that chromatin-accessibility evolves with developmental time (age) and follows along differentiation trajectories.

To validate the tree, we examined the gene accessibility of cell type-specific markers along the differentiation trajectory of individual lineages as well as compared it to a matching scRNA-seq tree. The scATAC-seq tree segregated known markers appropriately (Figure 3C). Moreover, the scATAC-seq tree had a comparable structure to that of a reduced scRNA-seq tree including only the three time points sampled for the scATAC-seq dataset (Supplementary information Figure 9B,C). Notably, putative near-projecting neurons (NP, Kim, Juavinett et al. 2015, Tasic, Yao et al. 2018) were the only population that was assigned to different branches in the scATAC-seq *vs*. scRNA-seq trees (Figure 3B *vs*. Supplementary information Figure 9B), probably reflecting the fact that these neurons may be molecularly related to both CFuPN and deep-layer CPN. This highlights the congruence of transcriptional and chromatin accessibility profiles at single-cell resolution from the developing cortex.

### Cis-regulatory cascades during differentiation of cortical cell types

In order to uncover changes in regulatory interactions during corticogenesis, we generated a map of all predicted cell type-specific *cis*-regulatory interactions by calculating co-accessible sites using the Cicero algorithm (Pliner, Packer et al. 2018) (Methods), identifying putative enhancers of known cell type-specific marker genes, and extracting the distal regulatory elements of these genes that are both active in and specific to each cell type (Figure 3D,E). As an example, we identified regulatory elements that are predicted to control *Pcp4,* a marker gene of SCPN and layer 6b neurons (Arlotta, Molyneaux et al. 2005, Molyneaux, Arlotta et al. 2005). In our transcriptional map and tree, *Pcp4* was also expressed in migrating neurons and was highly scored in the NMF gene program for neuronal differentiation and migration (Supplementary information Figure 5A). Several distal elements were co-accessible with the *Pcp4* putative promoter in the distinct cell types (Methods). Some were selectively co-accessible with the *Pcp4* gene in migrating neurons, suggesting developmental stage-specific enhancers (Figure 3F). Consistent with this interpretation, these enhancers contained binding sites for TFs associated with neuronal differentiation and migration, including *Nfix* and *Neurod2* (Figure 3F). Conversely, other *Pcp4* distal elements were selectively co-accessible in CFuPN, and contained binding sites for the SCPN identity regulators *Fezf2* and *Bcl11b*, suggesting a possible site for regulation of *Pcp4* in these neurons (Figure 3F). This approach also has the potential to identify novel regulators of cell fate. For example, predicted enhancers for *Pcp4* and *Fezf2* active in E18.5 SCPN had binding sites for the TF *Hivep2. Hivep2* was also predicted by our branchpoint analysis of the RNA tree (Figure 2E), where it was associated with CFuPN fates, possibly indicating a previously unknown role for this gene in CFuPN specification.

Next, we sought to identify putative TFs regulating distal elements active along development of individual lineages and at branchpoints. We searched for over-representation of known TF motifs in cell-type specific regulatory elements, both along the differentiation trajectories and in terminally differentiated cell types (Figure 3G,H). TF motifs were enriched along pseudotime cascades as differentiation progressed (Figure 3G), including sites for both TFs known to regulate cell identity and for novel ones. For instance, early segments of the cascade showed enrichment of *Dmrta2* motifs in AP-associated enhancers; a gene which has been recently associated with lissencephaly in humans(Urquhart, Beaman et al. 2016). In later stages, as neuronal cell types diverged, cascades included lineage-specific motifs, culminating in specific sets of TF binding sites enriched in the different cortical cell types (Figure 3H). For instance, *Cux1, Cux2,* and *Pou3f2* motifs were enriched in layers 2&3 CPN, consistent with their roles in CPN specification (Cubelos, Sebastian-Serrano et al. 2010). Motifs for *Bcl11b, Tbr1,* and *Fezf2* were enriched in CFuPN, together with less expected TFs such as *Nfe2l3* (Galazo, Emsley et al. 2016), *Nfia* (Plachez, Lindwall et al. 2008), and *Hivep2,* while astrocyte lineages were enriched for motifs for *Hes5, Sox9,* and *Klf3,* among others (Figure 3H).

The analysis of accessibility profiles thus identified the cis-regulatory elements and candidate regulatory factors defining differentiation and diversification of individual cortical cell types, further enhancing the inferred regulatory logic that drives lineage specification during cortical development.

### *Fezf2* controls the decision between CFuPN and CPN identity

Finally, we demonstrate how to use the framework of our single cell molecular atlas of normal mouse cortical development, as a new tool to elucidate phenotypic changes in loss-of-function models. We focused on the transcription factor *Fezf2* because its absence causes complete loss of SCPN (Chen, Schaevitz et al. 2005, Molyneaux, Arlotta et al. 2005, Lodato, Rouaux et al. 2011), consistent with it being one of the top-ranked genes in the SCPN differentiation program (Figure 4A). The mechanisms behind the loss of SCPN, and the identity of the neurons produced in their place, remain poorly understood.

We profiled 17,344 control (Het – heterozygous) and 16,117 knock-out (KO) cells by scRNA-seq from E15.5 and P1 developing cortex of *Fezf2* mutant mice(Hirata, Suda et al. 2004) (Supplementary information Figure 10A,B). To identify gene modules that varied between genotypes, we performed NMF gene module analysis on the combined *Fezf2* mutant and control data. The modules included all of the normal-development gene modules observed in the original E15.5 wild-type (WT) dataset (Supplementary information Figure 10C).

The modules corresponding to SCPN and CThPN specification were uniquely and highly downregulated in KO cells (Figure 4B). In our developmental atlas, *Fezf2* was a top-ranked gene in these modules, and was predicted as a candidate regulator of CFuPN lineages (Figure 2C-F and 4A). Moreover, ~70% of the 100 top-ranked genes in the SCPN and CThPN modules were downregulated in the mutant (*p*-value < 0.001, Wilcoxon Rank Sum test), suggesting that *Fezf2* is a regulator of the program, and not only a member gene (Figure 4B and Supplementary information Figure 10D).

Interestingly, two gene modules in the *Fezf2* mutant data did not match any E15.5 WT module. These KO-specific gene programs were enriched for Gene Ontology terms related to axon development and guidance (Figure 4B and Supplementary information Figure 10E), consistent with the fact that the mutant cells aberrantly project across the anterior commissure instead of subcortically (Lodato, Molyneaux et al. 2014). Overall, combining scRNA-seq profiles of a relatively small number of *Fezf2* mutant cells with the catalog of molecular programs from our comprehensive developmental atlas allowed us to successfully identify *Fezf2* as a key regulator of SCPN and CThPN specification programs, in an unsupervised approach.

Although prior investigation of *Fezf2* mutant mice had demonstrated that SCPNs were replaced by another neuron type, the exact subtype identity that these cells acquired remained elusive beyond a tentative association with deep-layer callosal projection neurons (Molyneaux, Arlotta et al. 2005, Lodato, Rouaux et al. 2011, Lodato, Molyneaux et al. 2014). This was largely because the full developmental trajectories of each neuronal cell type were not available, making it difficult to distinguish between the possibilities of an immature cell type *vs*. a specific terminal identity.

We thus applied our comprehensive corticogenesis atlas to define the class-specific identity of the aberrant cells emerging in the *Fezf2* mutant relative to normal cortical cell types. We first co-embedded the KO and control cells at both E15.5 and P1 (Figure 4C,D). In the KO, normal deep-layers neuron populations were lost, and replaced by a KO-specific deep-layers-like population at both ages (Figure 4C,D), indicating that in the absence of *Fezf2,* cells that would normally become SCPN or CThPN acquire aberrant identities not present in the control tissue.

To define the closest identity of these cells, we used the SingleCellNet method(Tan and Cahan 2019) to train a multi-class Random Forest classifier on the cell types of our developmental atlas, and then applied it to the KO-specific cells (Figure 4E and Supplementary information Figure 11A). The classifier assigned control cells with high accuracy, with errors confined to highly similar cells (*e.g*. layer 4 stellate neurons and CPN, Supplementary information Figure 11B). Interestingly, most of the KO cells were assigned into two groups: CThPN or layers 5&6 CPN identity (Figure 4E and Supplementary information Figure 11 B,C). While at E15.5, 22% of the KO-specific cells were classified as SCPN, at P1, only 1% of the KO-specific cells had this assignment (Figure 4E and Supplementary information Figure 11B,C). This suggests that a subset of these cells transiently express a rudimentary CFuPN/SCPN program, independent of *Fezf2.* However, virtually none of the KO-specific cells were assigned to migrating or immature identities at either age. Compared to endogenous CThPN, the KO-specific CThPN-like cells had higher expression of CPN genes, such as *Ptn, Serpine2,* and *Lpl* (Figure 4F). Relative to control deep-layers CPNs, the KO-specific CPN-like cells had higher expression of *Npy* and *Cck* and downregulation of *Nrgn,* among other differences (Supplementary information Figure 11D), and showed substantial divergence from control SCPN gene programs (Supplementary information Figure 11E). Consistent with the identities assigned by the classifier model, sub-clustering the KO-specific deep-layers neurons alone identified two subpopulations (Supplementary information Figure 11F-O), with the cluster partitioning matching the assignments made by the classifier.

Altogether, by comparison to our WT atlas of neocortical development, our analysis shows that loss of *Fezf2* alters the identity of CThPN, upregulating components of CPN gene programs, and results in the replacement of SCPN with cells resembling, but distinct from, layers 5&6 CPN (Figure 4G). This suggests that *Fezf2* has a key role in the suppression of CPN gene programs in developing corticofugal projection neuron populations. In addition, the aberrant populations do not represent cells stalled at any immature stage, but instead reach a terminal identity that differs from any endogenous neocortical cell type.

Lastly, we aimed to determine at which point in neuronal specification *Fezf2* acts. We profiled single cells from E13.5 control and *Fezf2* KO cortex by scRNA-seq, and did not find major differences in contribution by genotype or in gene expression in APs or IPs (Supplementary information Figure 12A-C). Only postmitotic neurons presented transcriptional differences, with a phenotype similar to that observed at the later time points (Supplementary information Figure 12D-E). Thus, although *Fezf2* is expressed in progenitors (Supplementary information Figure 4E, Guo, Eckler et al. 2013), its role in SCPN specification is post-mitotic. This supports our finding in the developmental atlas that neuronal subtype identity becomes restricted post-mitotically, and that a shared, potentially plastic transcriptional program governs the first steps of neuronal differentiation and specification.

## DISCUSSION

Extensive studies over the last ~20 years have helped identify some of the key genes that control the development of some of the main neuronal populations of the neocortex (Greig, Woodworth et al. 2013, Lodato and Arlotta 2015). However, the mechanistic principles by which the cerebral cortex generates its cellular diversity have remained elusive, because of the need to integrate all of its cell types, across all of their developmental stages, within a single framework. The advent of high-throughput single-cell molecular profiling has made it possible to explore developmental dynamics in heterogenous and complex tissues, such as very early areal specification in human cortex (Nowakowski, Bhaduri et al. 2017) and changes in apical progenitors as they contribute to the different cortical neuron populations (Yuzwa, Borrett et al. 2017, Telley, Agirman et al. 2019). We here present the first comprehensive single-cell expression atlas of neocortical development, profiled at high temporal resolution to generate a description of all of its cell types and states. The data expand our knowledge of the principles that shape cell diversification during corticogenesis.

It is still debated when during the development of neuronal lineages cellular diversification occurs, such as the possible contribution of progenitor heterogeneity to generating the diverse projection neuron populations. Our data supports the model that progenitors gradually change in state over the period of excitatory neurogenesis (Yuzwa, Borrett et al. 2017, Telley, Agirman et al. 2019), rather than having separate lineage-restricted transcriptional programs (Franco, Gil-Sanz et al. 2012). We find that the transcriptional programs of excitatory lineages emerge from a common progenitor branch and diverge gradually over the course of post-mitotic development. We did not find clear distinctions among progenitors that would support a model of discrete, strictly fate-restricted, progenitor populations, nor did we observe separate trajectories linking progenitors of different ages with projection neurons of distinct layers. Transcriptional programs of excitatory lineages diverged only in post-mitotic neurons, and broad molecular programs were widely shared by PN subtypes throughout corticogenesis.

The availability in the field of a wide range of single-gene mutant mice presents an invaluable resource to help understand how specific events of corticogenesis are controlled. Our comprehensive molecular atlas of all cortical lineages represents a novel tool for phenotyping mutants without prior knowledge of gene function. Using the *Fezf2* mutant as a test case, we determined that *Fezf2* controls gene programs driving identity and specification of both SCPN and CThPN. It is interesting that the cells produced in place of SCPNs in the mutant are similar to but also clearly distinct from layers 5&6 CPNs. We speculate that these aberrant cells may reflect an ancestral commissural neuronal identity, given that these cells project to the anterior commissure (Lodato, Molyneaux et al. 2014), a structure that is prominent in species that lack the corpus callosum, such as reptiles.

Determining the chromatin accessibility landscape across lineage trajectories enabled us to define cell type-specific enhancers and candidate transcriptional regulators of PN specification. Interestingly, layer 5&6 CPN share enhancer elements with CFuPN, which have a similar birthdate (Figure 3D); for instance, the *Sox5* motif was enriched in both deep-layers CPN and CFuPN. This may reflect a “birthdate memory” or broadly-shared layer identity. Consistent with this, loss of *Fezf2* caused SCPN populations to acquire similarities to deep-layers CPN, perhaps indicating an epigenetic landscape that is receptive for that fate. Our lineage-tree reconstruction uncovered expression of genes that are typically considered specific to individual cell types in other populations. For instance, *Pcp4*, which is traditionally considered a CFuPN marker, is also transiently expressed in other neuron types as they migrate to the cortical plate. Correspondingly, in our chromatin accessibility analysis we were able to identify separate *Pcp4* enhancers predicted to be active specifically in migrating neurons and in corticofugal projection neurons. These findings suggest novel roles for genes traditionally considered markers of single lineages.

This work provides the first comprehensive collection of all of the molecular states of each cortical lineage through time, and help identify candidate molecular effectors and regulatory elements underlying fate divergence. This information can be directly applied to functional interrogation of candidate gene programs and regulators using scalable genetic assays, such as Perturb-seq (Dixit, Parnas et al. 2016), and mutant models to ultimately parse the core mechanisms that drive development of the neocortex.

**Figure 1.**
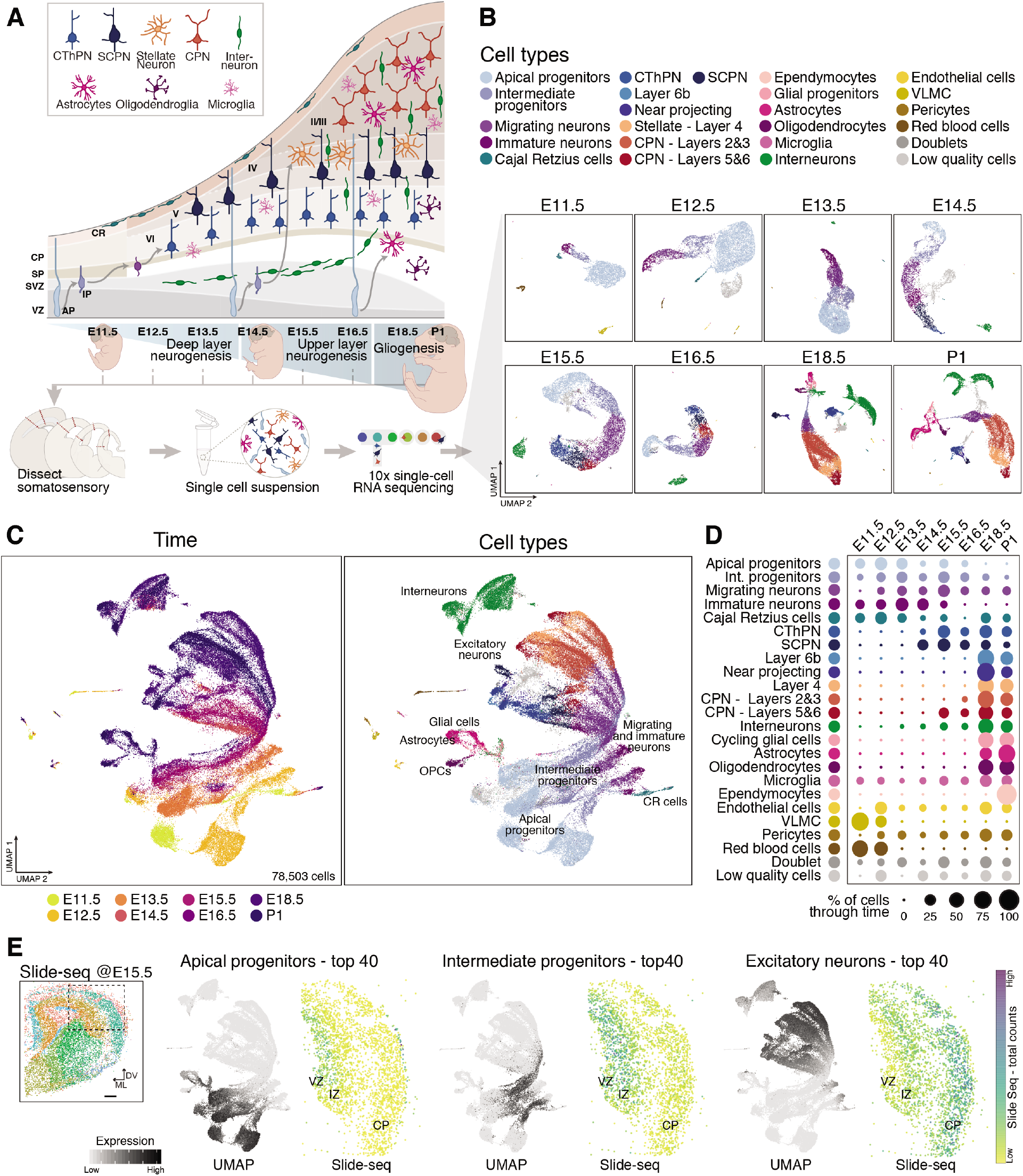
Comprehensive atlas of murine cortical development. **A** Schematic showing the cellular diversity and sequential generation of projection neuron subtypes and glial cells in the neocortex, as well as the experimental approach and time points collected to build the developmental atlas. **B** Visualization of scRNA-seq data collected at the different time points by UMAP. Cells are colored according to their cell types. Cell types were assigned by expression of marker genes and evaluation of differentially expressed genes. **C** Combined time points plotted together and visualized by embryonic stage of collection (left), or cell types (right), legend in **D**. **D** Distribution of each cell type across the different embryonic ages. **E** Combined expression of the top 40 differentially expressed genes for the indicated cell types overlaid on the scRNAseq data (to the left for each cell type) or in coronal sections of an E15.5 brain processed by Slide-seq (to the right for each cell type), where each dot represents a bead colored according to the number of counts. Far left: the same section, with dots colored by their predicted cell identity. Boxed region corresponds to area shown in right panels. VZ: ventricular zone, SVZ: subventricular zone, SP: subplate, CP: cortical plate, CR: Cajal Retzius cells, AP: apical progenitors, IP: intermediate progenitors.

**Figure 2.**
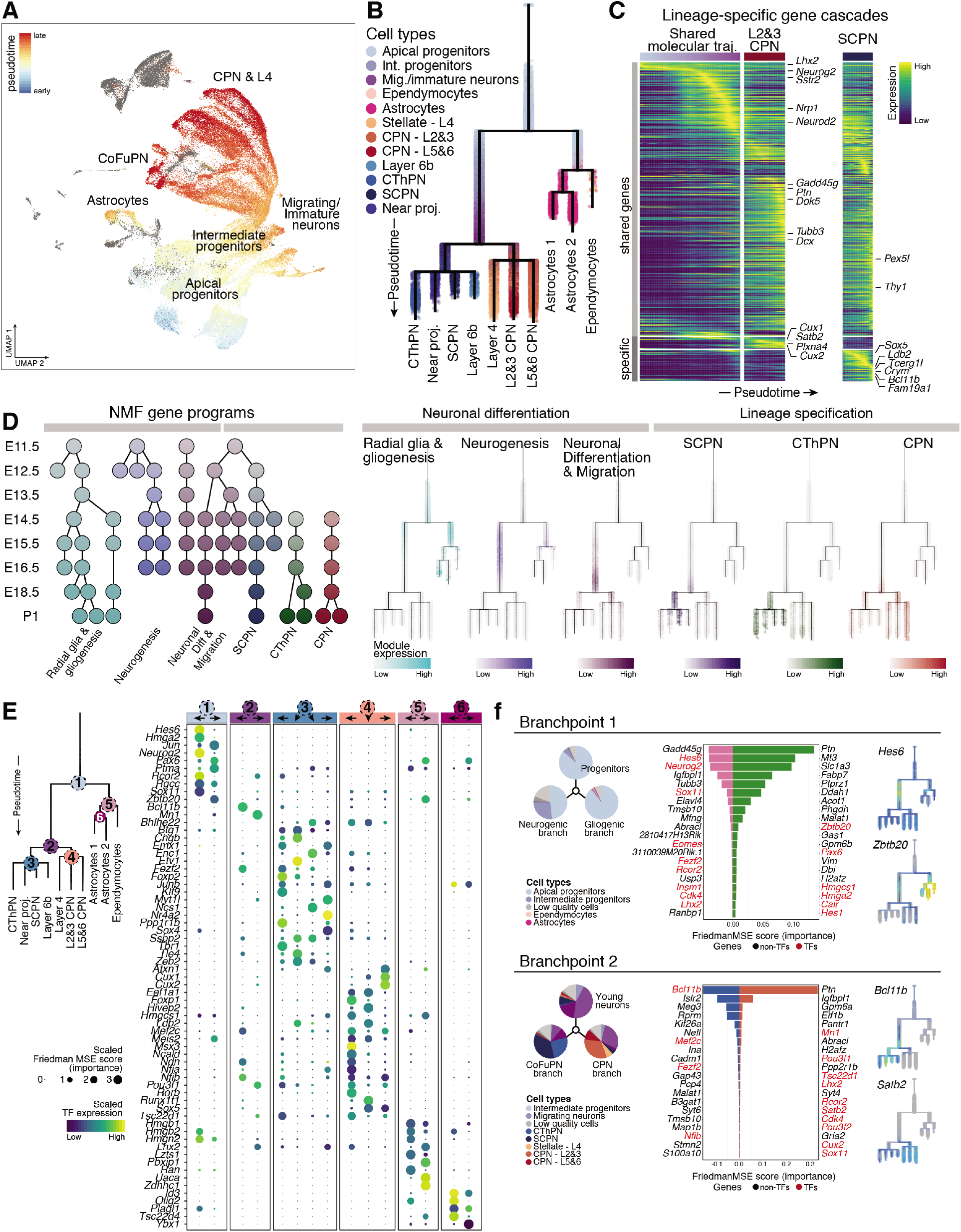
Molecular developmental trajectories of neural cell types in the neocortex. **A** UMAP visualization of the scRNA-seq data from combined time points, with cells colored by pseudotime. Blue represents earlier cells in the trajectory, while red labels later cells. Grey: cells that were excluded from the trajectory. **B** URD trajectory branching tree of the developing cortex. Cells are organized based on their pseudotime value and their transcriptional similarity. Root is earliest progenitors at E11.5, tips are terminal cell types collected at P1. Cells are colored according to their identity (as in Figure 1C). **C** Smoothened heatmap of gene cascades for layers 2&3 CPN and SCPN differentiation. The x axis represents pseudotime across the tree. Each row is a gene where gene expression is scaled to the maximum observed expression, and then smoothened. Marker genes are labeled, and neuronal sub-type specific genes are grouped at the bottom. The cascade is divided into three segments: the shared trajectory between both cell types (left), layers 2&3 CPN-specific (middle) and SCPN-specific (right) trajectories. Cascades for all trajectories (with all genes labeled) are presented in Supplementary information Figs. 6–8. **D** Gene programs of connected modules found by NMF. Left: Each circular node represents a module, horizontally aligned to the developmental stage they were computed from and colored by their identity. Right: Scaled expression of modules corresponding to individual programs, colored according to program identity. **E** Feature importance and average expression of genes predicted to be involved in cell types divergence. Top 5 genes per branch, ranked by their Friedman MSE score (importance) for distinguishing between daughter branches. Color bar at top indicates branchpoints marked on the tree to the left. Arrows indicate daughter branches. **F** For branchpoints 1 and 2, pie charts showing cell-type distribution immediately before and after branch divergence (left). Middle: Bar plots show importance score in left- or right-facing branches. Transcription factors are labelled in red. Right: Examples of gene expression for genes distinguishing between trajectories.

**Figure 3.**
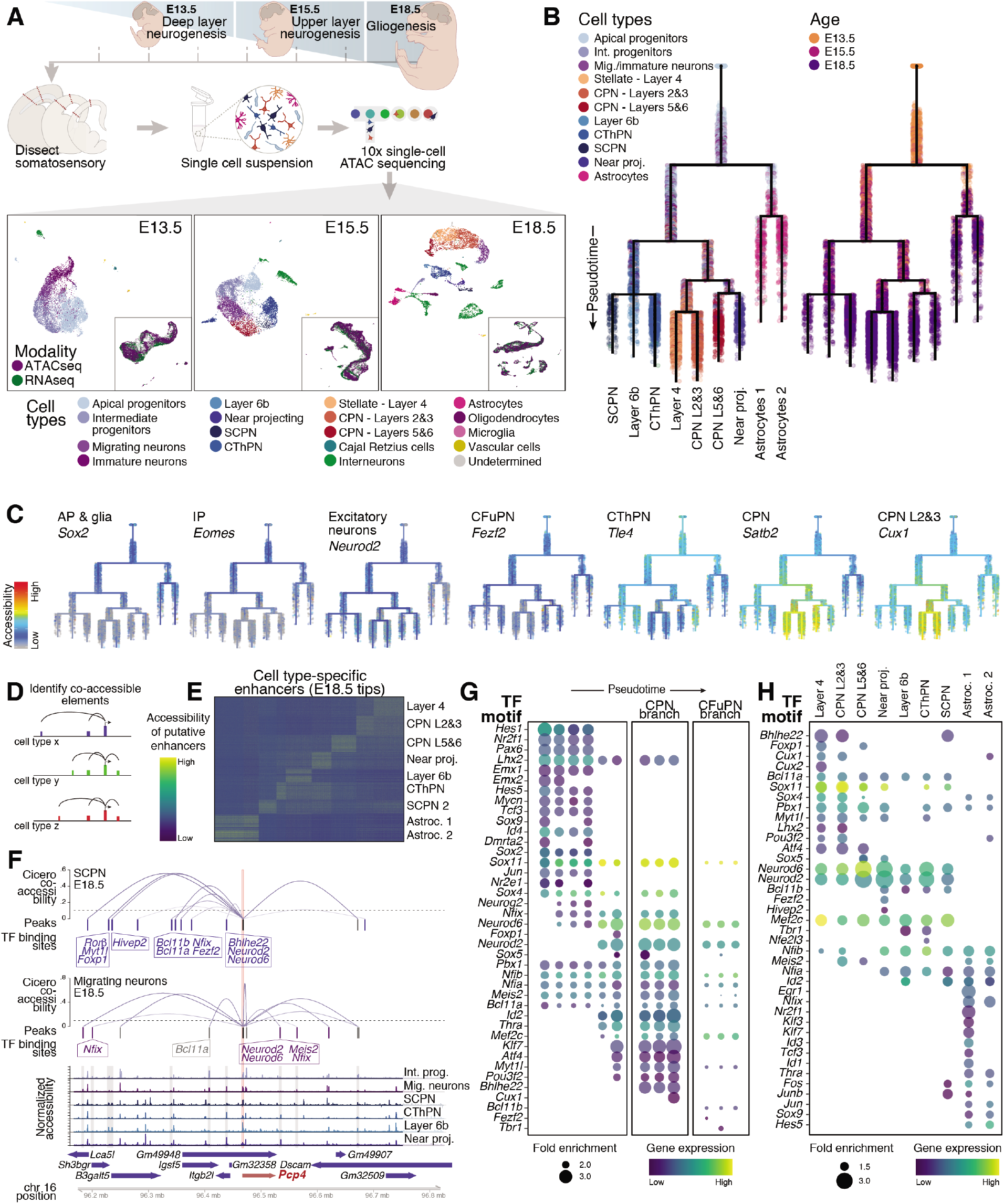
scATAC-seq landscape of the developing neocortex reveals cell type-specific enhancers throughout development. **A** Schematic showing the experimental approach and time points collected for scATAC-seq. UMAP visualization of the accessibility data for each time point. Cells are colored by their predicted cell type from integration with corresponding scRNA-seq datasets. All cell types recovered from transcriptomes were also identified by scATAC-seq. Insets show consistent integration of RNA and ATAC data colored by modality. **B** URD chromatin accessibility trajectories during cortical development. Cells are organized based on pseudotime values. Root is earliest progenitors at E13.5, tips are final cell types obtained at E18.5 (with an identity-prediction score higher than 70%). Cells are colored according to their identity as predicted from scRNA-seq (left) or age of collection (right). **C** ATAC trees highlighting the accessibility of marker genes characteristic of the different cortical cell types, including apical and intermediate progenitors, excitatory neurons, callosal neurons, layer 4 stellate neurons and corticofugal neurons. **D** Schematic of the approach used to identify candidate cell type-specific enhancers. Differential expression analysis identified cell type-specific genes, for which we calculated co-accessibility (correlation higher than 25%) between distal elements (within a 250 kb region) and target gene promoters using Cicero, within each cell type. **E** Candidate cell type-specific enhancer regions identified at E18.5 for cells derived from dorsal progenitors, as indicated in **D**. **F** Distal elements co-accessible with the *Pcp4* promoter region in E18.5 SCPN and migrating neurons. Cicero co-accessibility is shown in blue curves, detected peaks in each cell type are shown as colored bars. Black bars correspond to promoter peak, blue bars are peaks selectively co-accessible in CFuPN, and purple bars are peaks only co-accessible in migrating neurons. Boxes indicate transcription factors whose motifs are present in indicated peaks. Peaks are aligned to coverage plots (bottom) showing combined ATAC reads for the indicated cell types. Chromosome coordinates and genes are indicated at bottom. **G-H** Transcription factor motifs enriched along the ATAC tree (**G**). The dot size shows fold enrichment, and the color is average RNA expression in nearest cells in the integrated RNA and ATAC data. Motif enrichment was calculated for each segment of the tree and separation between columns indicates branchpoints. Motif enrichment for the tree tips (terminal cell types) is shown in **H**.

**Figure 4.**
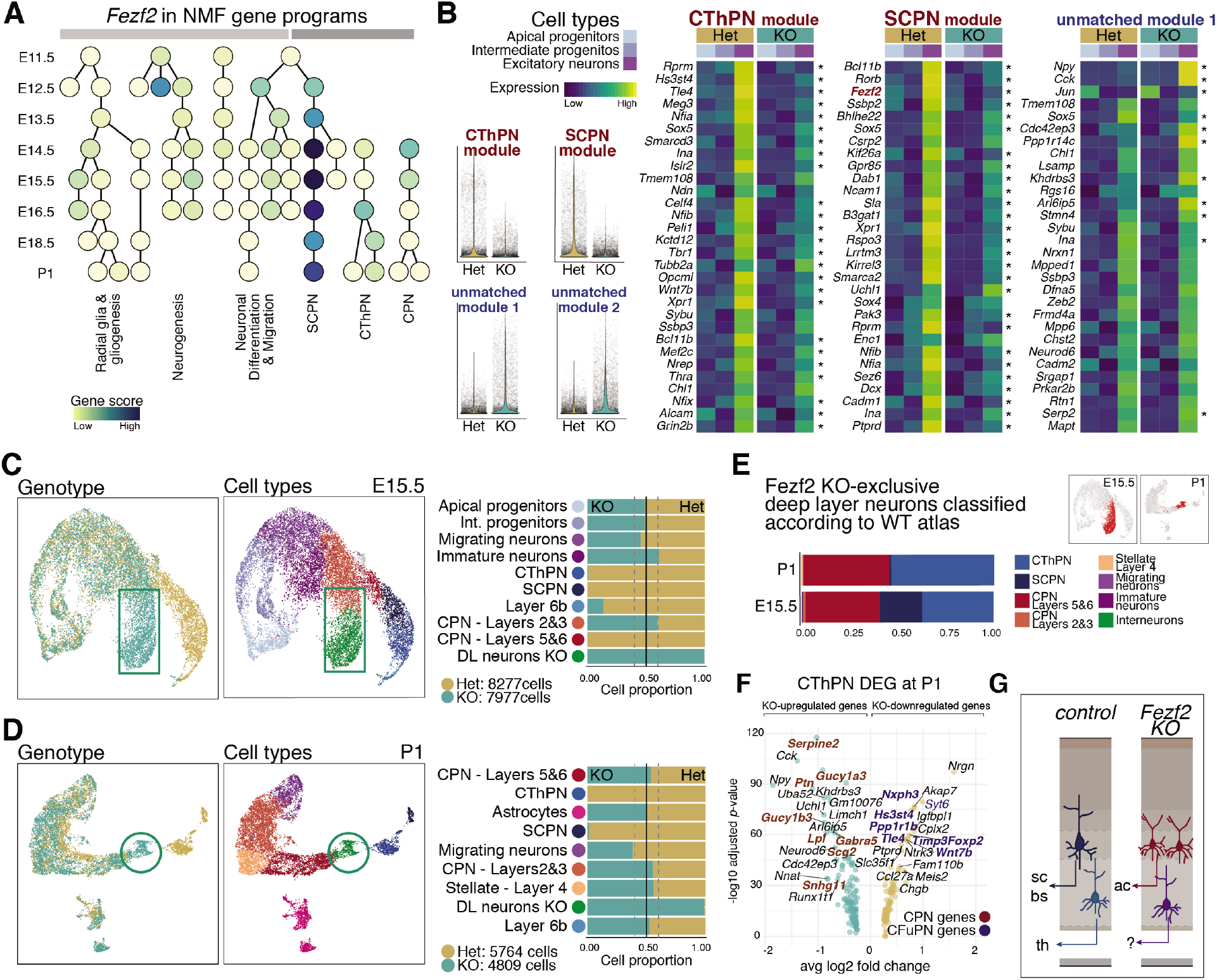
Fezf2 prevents acquisition of callosal identity in CFuPNs. **A** Gene programs of connected modules, as in Figure 2D, colored by *Fezf2* participation (score) in each module. **B** Expression of affected modules in *Fezf2* KO E15.5 cortex. Left: Violin plots showing scaled module expression in individual cells. Only affected modules are shown. Right: Heatmap showing average expression of the top 30 genes for selected modules, in three major cell types, apical and intermediate progenitors and excitatory neurons, by genotype. Asterisk indicates whether a gene is differentially expressed between control and KO neurons, at the single cell level (adjusted *p*-value < 0.001, Wilcoxon Rank Sum test with Bonferroni correction for multiple testing). **C-D** UMAP visualization of single-cell transcriptomes collected from control (*Fezf2* heterozygous) and KO cortices at E15.5 (**C**) and P1 (**D**), colored by genotype (left) and cell type (center). Only dorsally-derived cells are plotted. Also shown is proportion of cells in each cell type by genotype (left). **E** Cell-type assignment of the cells in the KO-specific cell clusters (UMAP insets, highlighted in red) by a classifier trained on the wild-type developmental time course. **F** Differential expression analysis of CThPN control cells and aberrant CThPN KO cells shows upregulation of CPN marker genes (in red) and downregulation of CThPN markers (blue). **G** Summary of phenotype in *Fezf2* KO. CThPN acquire an aberrant phenotype and upregulate CPN genes, while SCPN convert to layers 5&6 CPN-like neurons and project through the anterior commissure (ac) to the contralateral cortex. th: thalamus, sc: spinal cord, bs: brain stem.

## SUPPLEMENTARY INFORMATION

**Supplementary information Figure 1 (related to Figure 1).**
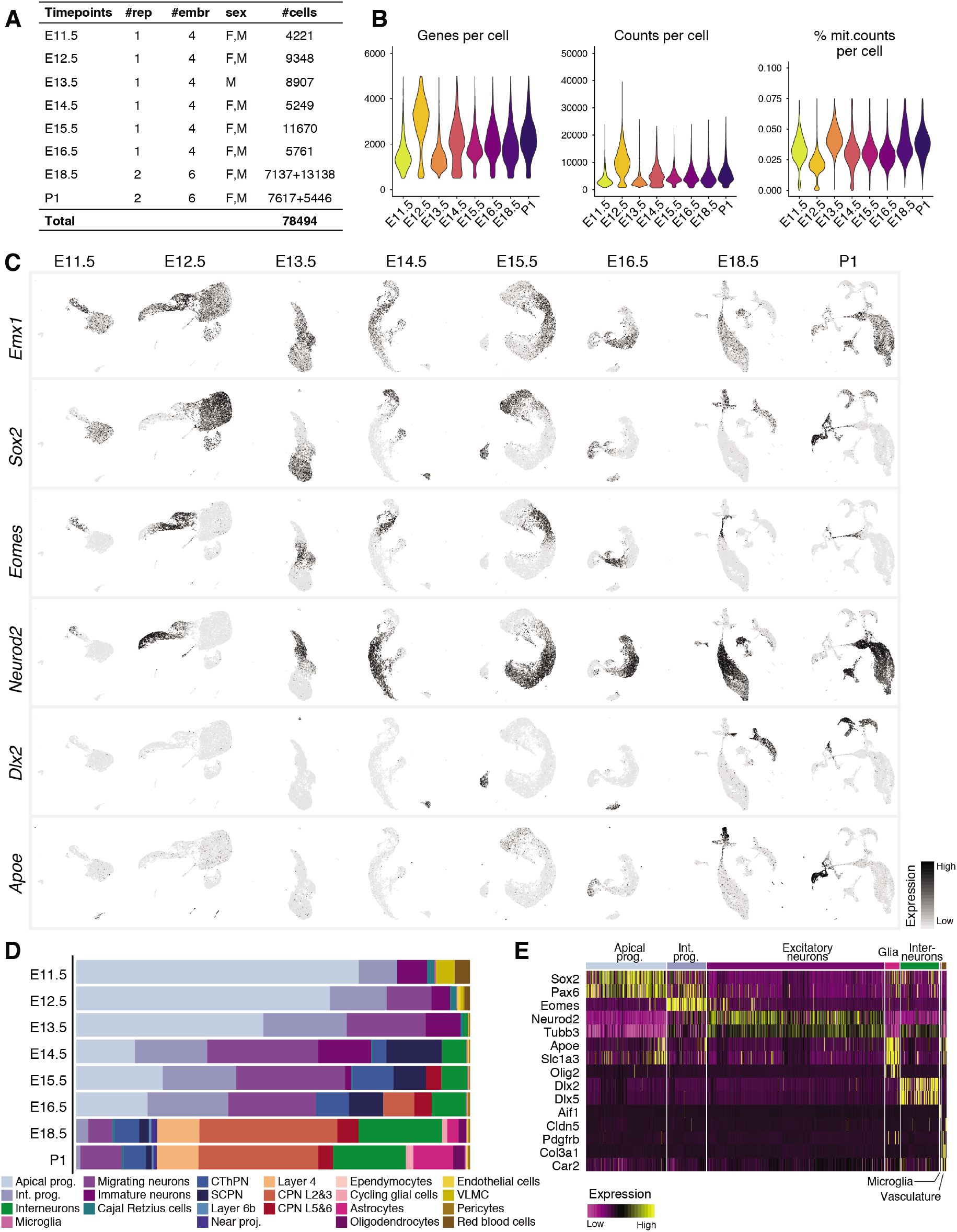
Classification of cell types in scRNA-seq data from individual embryonic ages. **A** Number of replicates, total number of embryos, sex of animals and number of cells analyzed per time point. **B** Number of genes, number of mRNA molecules, and percentage of mitochondrial counts per cell in each time point. **C** UMAP visualization of cells collected at each time point, showing expression levels of marker genes for dorsal derivatives *(Emx1),* apical progenitors *(Sox2),* intermediate progenitors (*Eomes*), excitatory neurons (*Neurod2*), inhibitory interneurons (*Dlx2*), and glial cells (*Apoe*). **D** Proportion of cells corresponding to the different cell types present in each time point. 85 to 98% of cells were successfully identified for each time point. **E** Expression of marker genes for the main cell types in the combined dataset. Each column is a cell and cells are grouped by cell type.

**Supplementary information Figure 2 (related to Figure 1).**
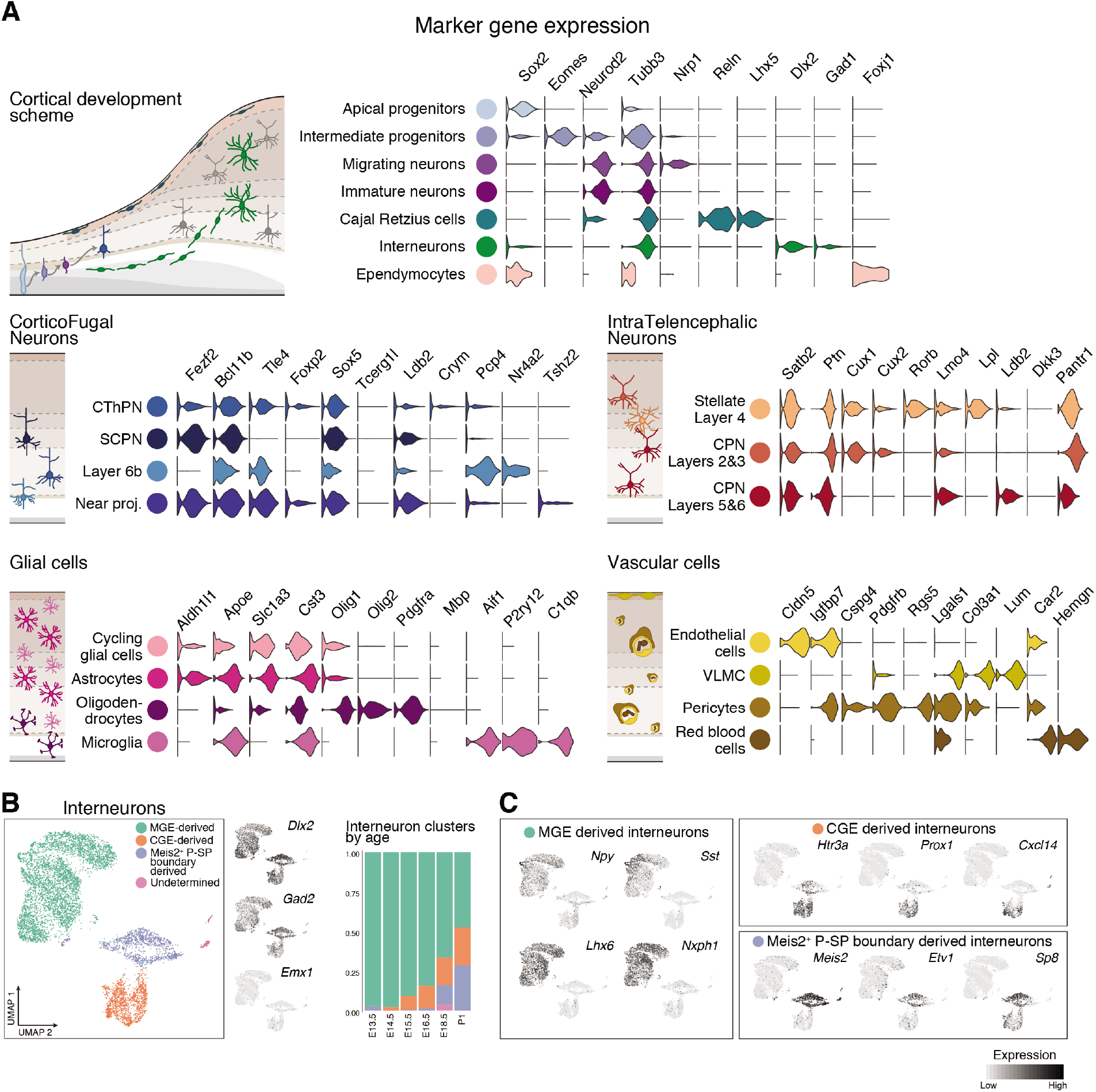
Molecular characterization of cell types in the developing cerebral cortex. **A** Selective expression of marker genes per cell type in the combined scRNA-seq dataset. Cell types are grouped based on their identity and shared marker genes. Far left: schematic of development of the cell types presented. **B** Different subtypes of interneurons integrate into the developing cortex through time. From left to right: clustering of interneurons collected at all time points, visualized via UMAP. Interneuron UMAP plots show the expression of the inhibitory markers *Dlx2* and *Gad2,* as well as a marker of dorsally-derived cell types (*Emx1*), not expressed by interneurons. Proportion of cells corresponding to each cluster in each time point. **C** Expression of genes characteristic of interneurons of different embryonic origins. MGE-derived interneurons express *Npy, Sst, Lhx6* and *Nxph1.* Interneurons originating in the CGE are positive for *Htr3a, Prox1, Cxcl14* and *Sp8.* A second population of *Htr3a^+^* interneurons express *Meis2, Etv1* and *Sp8*.

**Supplementary information Figure 3 (related to Figure 1).**
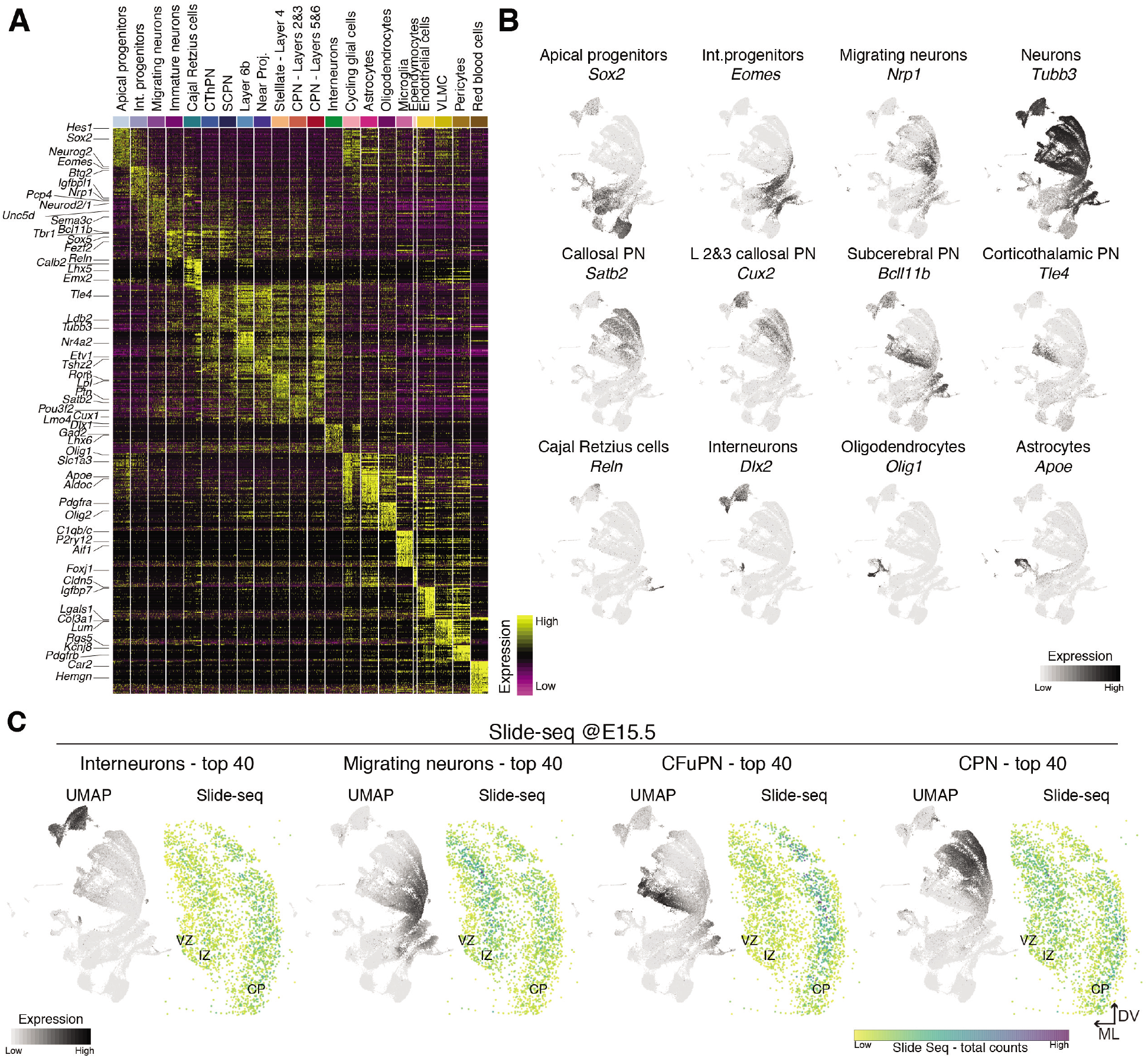
Molecular signatures and spatial distribution of cell types in developing cerebral cortex. **A** Gene signatures for all cell types identified in the combined time points. Top 20 differentially expressed genes for each cell type are presented. Cells were down-sampled to a maximum of 500 cells per cell type. **B** Expression of canonical marker genes for selected cells in the scRNA-seq time course. **C** Combined expression of the top 40 differentially expressed genes for the indicated cell types overlaid on the scRNA-seq data (to the left for each cell type, scores calculated by AddModuleScore from Seurat) or in coronal sections of an E15.5 brain processed for Slide-seq (to the right for each cell type), where each dot represents a bead colored according to the total number of counts.

**Supplementary information Figure 4 (related to Figure 2).**
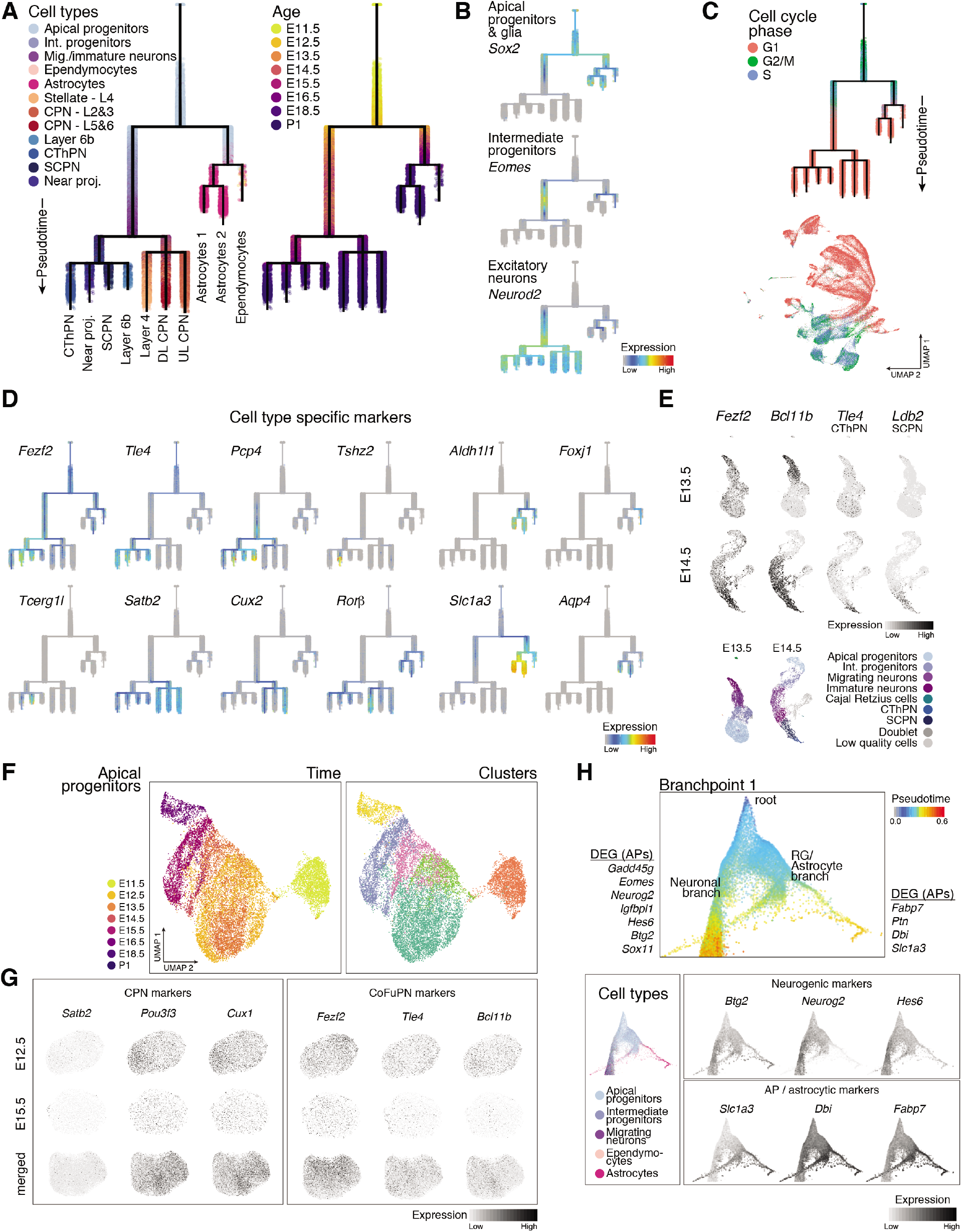
Neuronal cell types diverge post-mitotically. **A** URD trajectory branching tree of the developing cortex. Cells are colored according to their identity (left) or developmental time of collection (right). **B** Branching trees labelled by the expression of marker genes *Sox2*, *Eomes* and *Neurod2*, showing the distribution and sequential developmental progression of apical progenitors, intermediate progenitors and excitatory neurons, respectively. **C** Branching tree and UMAP representation of full developmental atlas colored by cell cycle phase as predicted by gene expression. Separation in different classes of neurons occurs post-mitotically. **D** Branching trees showing the expression of marker genes characteristic of the dorsally-derived cortical cell types, including callosal neurons *(Satb2, Cux2),* layer 4 stellate neurons *(Rorb),* corticofugal neurons *(Fezf2, Tle4, Pcp4, Tcerg1l),* putative near-projecting neurons *(Tshz2),* astrocytes *(Slc1a3, Aqp4, Aldh1l1),* and ependymocytes *(Foxj1).* **E** Expression of CFuPN marker genes at E13.5 and E14.5. Newborn neurons at E13.5 do not express *Tle4* or *Ldb2,* markers of CThPN and SCPN, respectively. By E14.5 *Tle4* and *Ldb2* become expressed in different clusters, indicating that CThPN and SCPN become distinguishable. Bottom panel, UMAP visualizations colored according to cell type are included for reference. **F** Apical progenitors from different ages form a continuous progression of cells and do not segregate into distinct clusters. Apical progenitors from all time points were sub-clustered separately, colored by age and clusters identified by Seurat. **G** Expression of CPN markers (*Satb2*, *Pou3f3* and *Cux1*, left), and CFuPN markers (*Fezf2*, *Tle4* and *Bcl11b*, right) in both early (E12.5) and late (E15.5) APs, as well as in the combined AP populations. APs were coembedded using the top 100 differentially expressed genes between CFuPN and CPN) as input for principal component analysis and downstream clustering and visualization. Marker genes are expressed in progenitors but do not drive clustering of the cells. **H** Forced layout embedding representation of the developmental branching tree, showing the initial part of the tree. Cells are colored according to their pseudotime value. The tree is labelled with the identities of the corresponding branches, and differentially expressed genes between APs in each branch are shown. Bottom panels, cells were colored by cell type (left) or according to expression levels of differentially expressed genes. Cells corresponding to the astrocytic and neuronal branches form a continuum of cells.

**Supplementary information Figure 5 (related to Figure 2).**
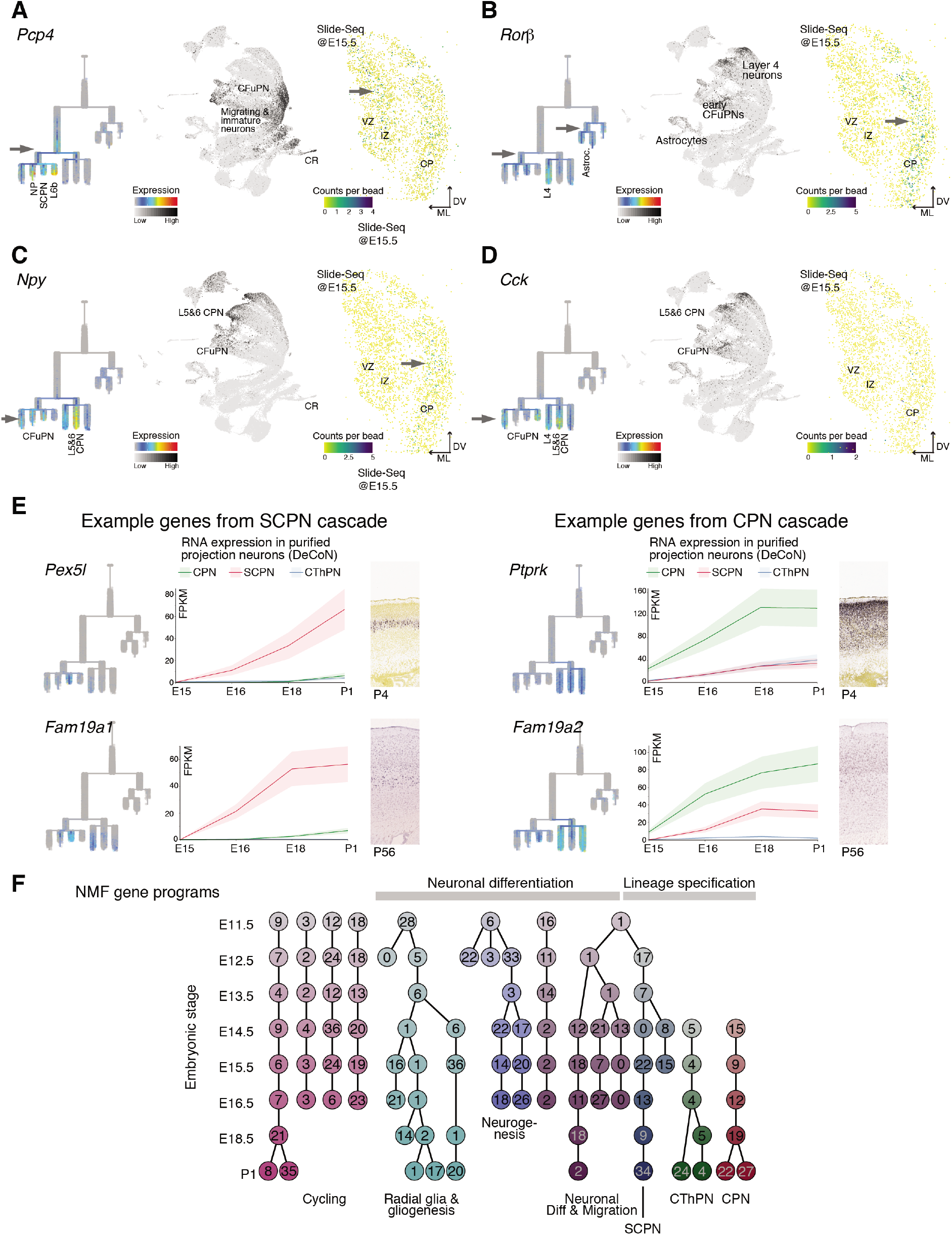
Novel expression pattern of selected genes. **A-D** Novel expression patterns emerging from the inferred tree. Expression levels overlaid on the tree diagram (left) and UMAP of full scRNA-seq developmental data (middle), and Slide-seq counts on an E15.5 section of cortex (right) for each gene. *Pcp4* is expressed in migrating and immature neurons that contribute to both CPN and CFuPN, as well as in SCPN, layer 6b, NP and CR cells, and is found in the intermediate zone (IZ) and cortical plate (CP) (**A**). *Rorb* is expressed in developing CFuPN, astrocytes and layer 4 stellate neurons and present in the deep CP (**B**)*. Npy* is expressed in CFuPN and highly in CPN of layers 5&6. Positive *Npy* signal is evident in the deep CP through Slide-seq (**C**). *Cck* was also detected in CFuPN and at higher levels in CPN of layers 5&6. Low levels of expression in the CP were detected via Slide-seq (**D**). VZ: ventricular zone. **E** Validation of expression of novel cell type-specific genes emerging from the cascade analysis. Expression levels overlaid on the tree diagram (left), time course expression on purified subtypes of PN from DeCoN transcriptomic resource (Molyneaux, Goff et al. 2015) (middle), and *in situ* hybridization from the Allen Developing Mouse Brain Atlas (right, age indicated in figure). **F** Complete set of gene programs of connected modules found by NMF. Each circular node represents a module. Modules are horizontally aligned to the developmental stage the module was computed from, and colored by similar program function or identity.

**Supplementary information Figure 6 (related to Figure 2).**
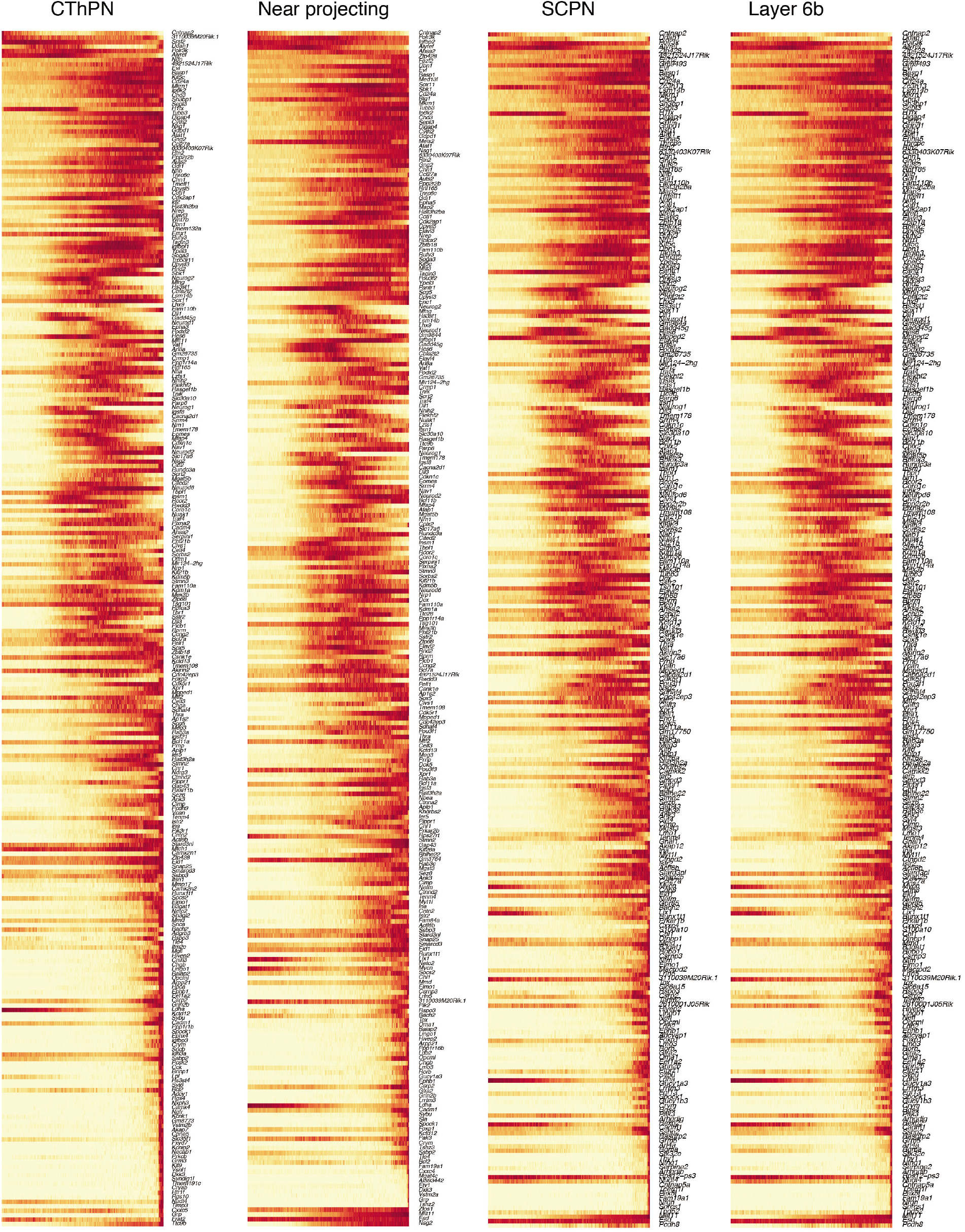
Genetic cascades accompanying development of corticofugal projection neurons. Gene cascades for CThPN, putative near projecting neurons, SCPN and layer 6b neurons differentiation (in that order from left to right). The x axis represents pseudotime across the tree. Each row is a gene where gene expression is scaled to the maximum observed expression, and then smoothened. All genes are labeled.

**Supplementary information Figure 7 (related to Figure 2).**
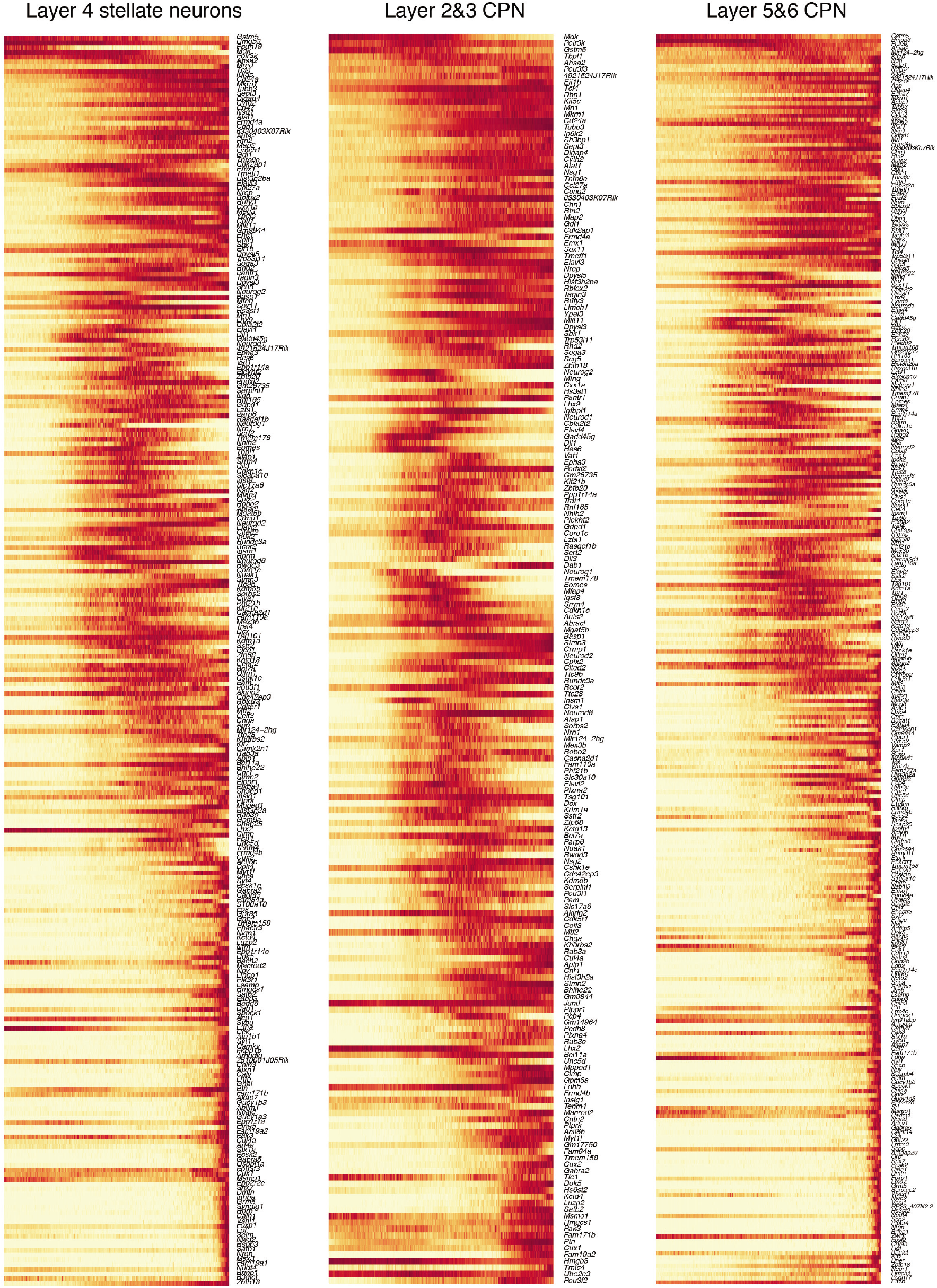
Genetic cascades accompanying development of callosal and and layer 4 neurons. Gene cascades for layers 2&3 and layers 5&6 (left and middle) and layer 4 stellate neuron (right) differentiation. The x axis represents pseudotime across the tree. Each row is a gene, where gene expression is scaled to the maximum observed expression, and then smoothened. All genes are labeled.

**Supplementary information Figure 8 (related to Figure 2).**
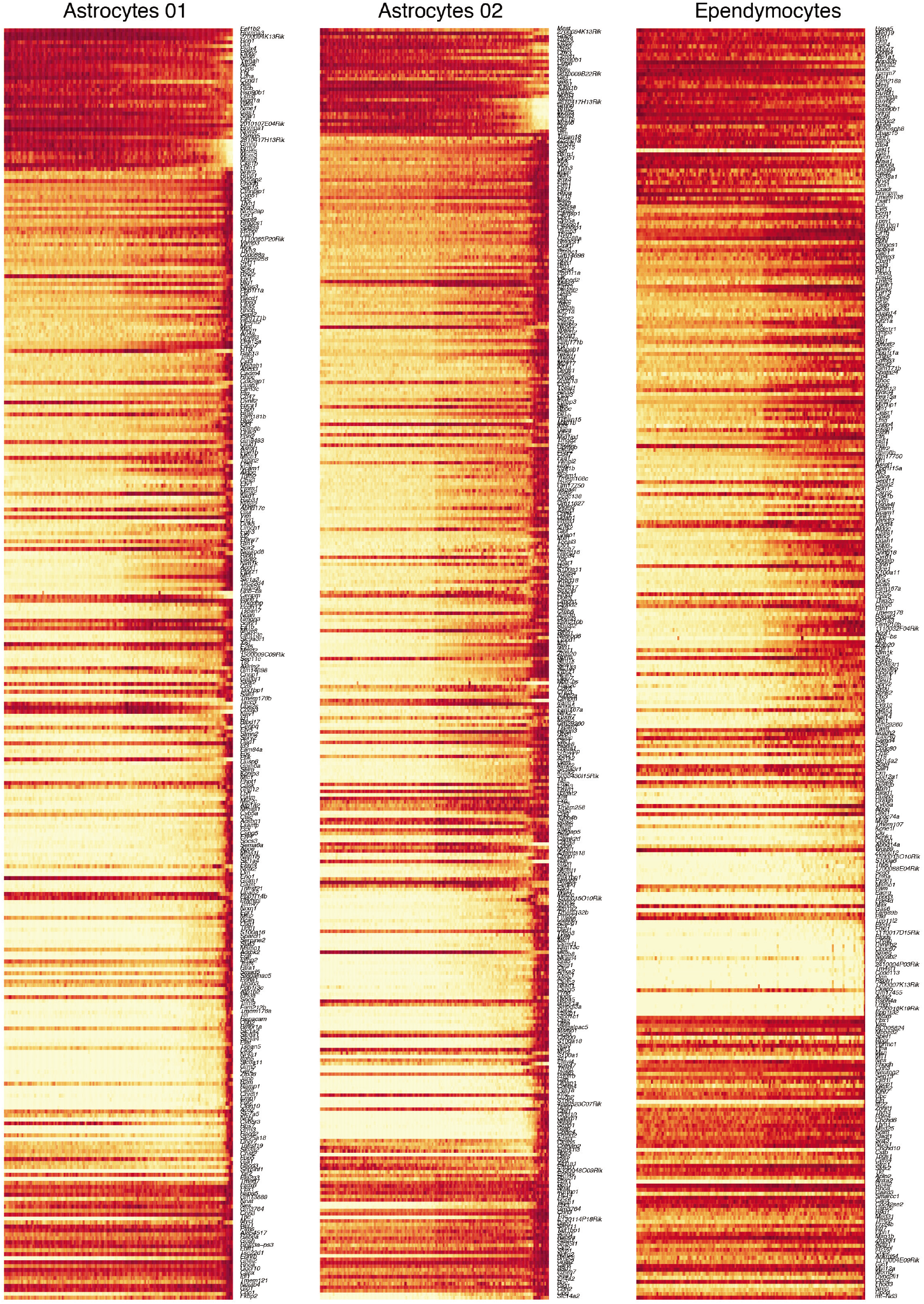
Genetic cascades accompanying development of astrocytes and ependymocytes. Gene cascades for astrocyte (left and middle) and ependymocyte (right) differentiation. The x axis represents pseudotime across the tree. Each row is a gene where gene expression is scaled to the maximum observed expression, and then smoothened. All genes are labeled.

**Supplementary information Figure 9 (related to Figure 3).**
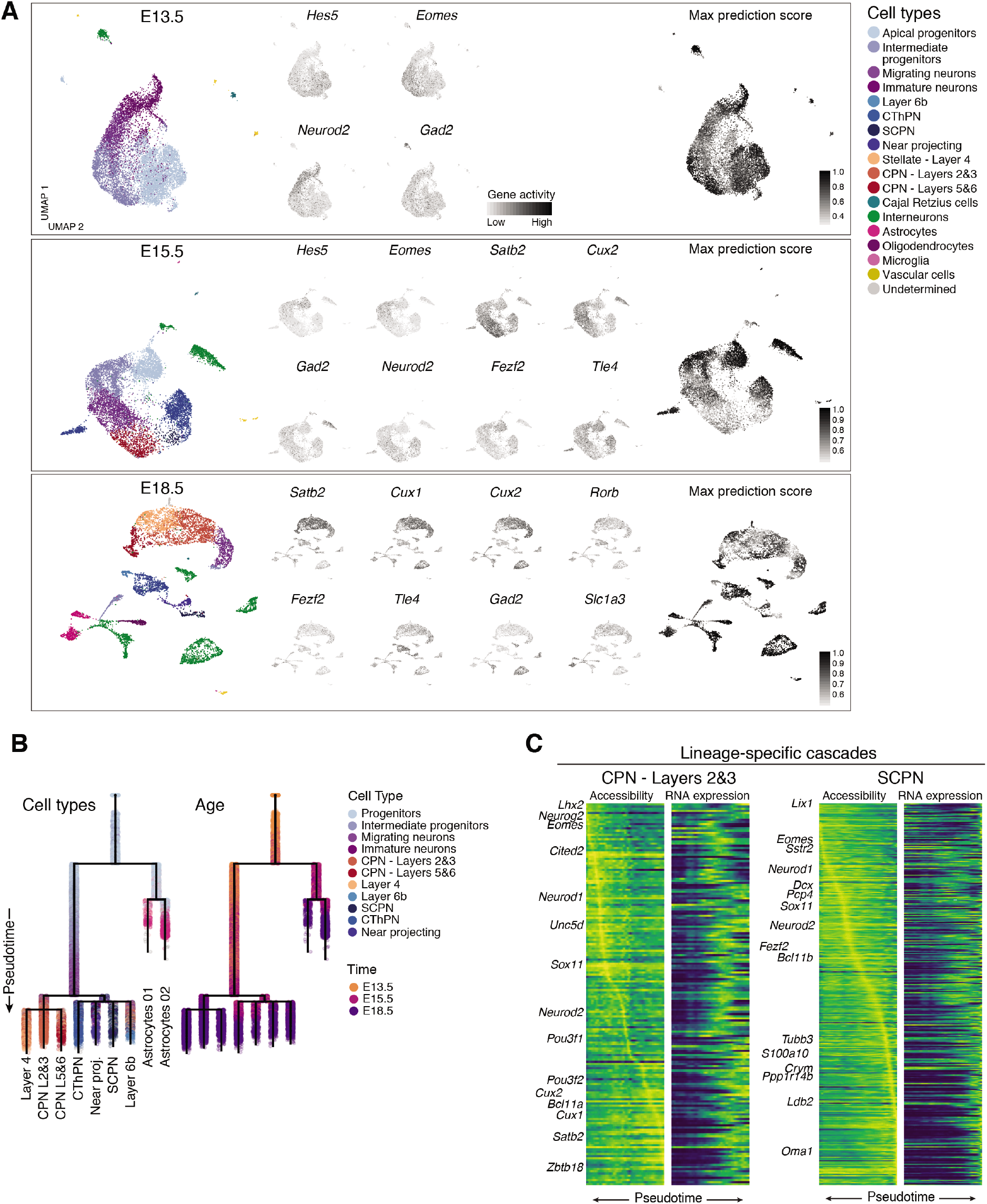
Characterization of a scATAC-seq atlas of cortical development. **A** scATAC-seq data per time point. UMAP visualization of the single cells colored by their predicted identity from integration with scRNA-seq datasets (left). Gene accessibility of selected markers for main cell types present in each time point (middle). Maximum prediction score for each cell based on labels transferred from scRNA-seq data (right). **B** RNA-based tree generated from only the E13.5, E15.5 and E18.5 time points, corresponding to the scATAC-seq data. Trees are colored by cell type (left) and time of collection (right). **C** Chromatin accessibility and gene expression cascades for layers 2&3 CPN and SCPN. Same genes are plotted for both modalities.

**Supplementary information Figure 10 (related to Figure 4).**
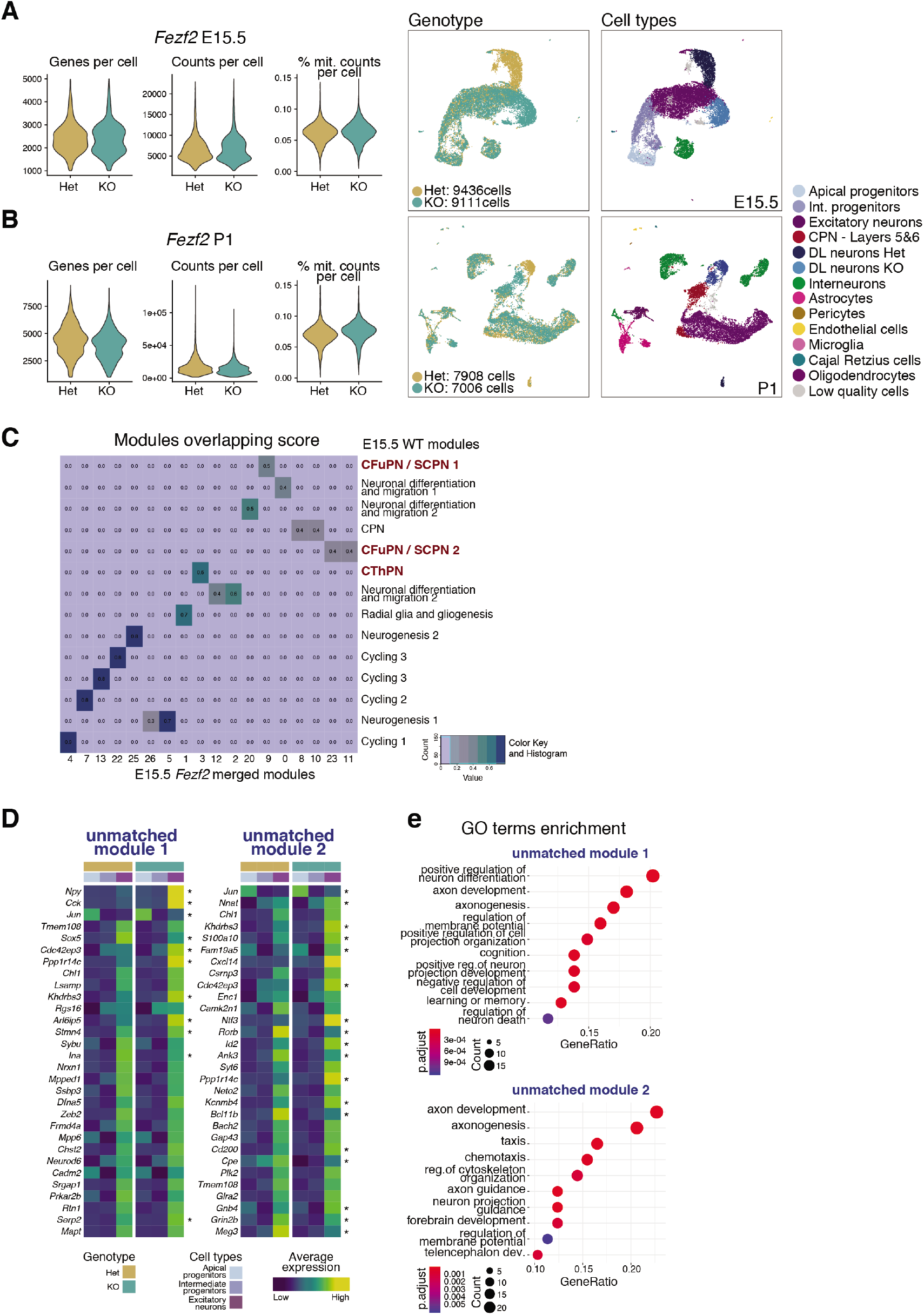
scRNA-seq analysis of *Fezf2* knock-out cerebral cortex. **A-B** Violin plots of number of genes (left) and mRNA molecules (counts; middle), percentage of mitochondrial counts (right) per cell in control (Het) and KO *Fezf2,* and UMAP visualizations of merged scRNA-seq data sets at E15.5 (**A**) and P1 (**B**). UMAP visualizations are colored by genotype or assigned cell type. **C** A heatmap showing the overlapping scores between NMF modules identified in the E15.5 *Fezf2* datasets and the original E15.5 wild-type modules. All modules were identified with an overlapping score of 40% or higher. **D** Expression of the top 30 genes for modules that showed differential levels between KO and control (Het) cells. Average expression was calculated for three mayor cell types, apical and intermediate progenitors and excitatory neurons, by genotype. The asterisk indicates whether a gene is differentially expressed between control and KO neurons, at the single-cell level (adjusted *p*-value < 0.001, Wilcoxon Rank Sum test with Bonferroni correction for multiple testing). Bottom, module “expression” levels overlaid on *Fezf2* E15.5 KO and control cells). **E** Gene ontology terms enriched in *Fezf2* KO-specific modules.

**Supplementary information Figure 11 (related to Figure 4).**
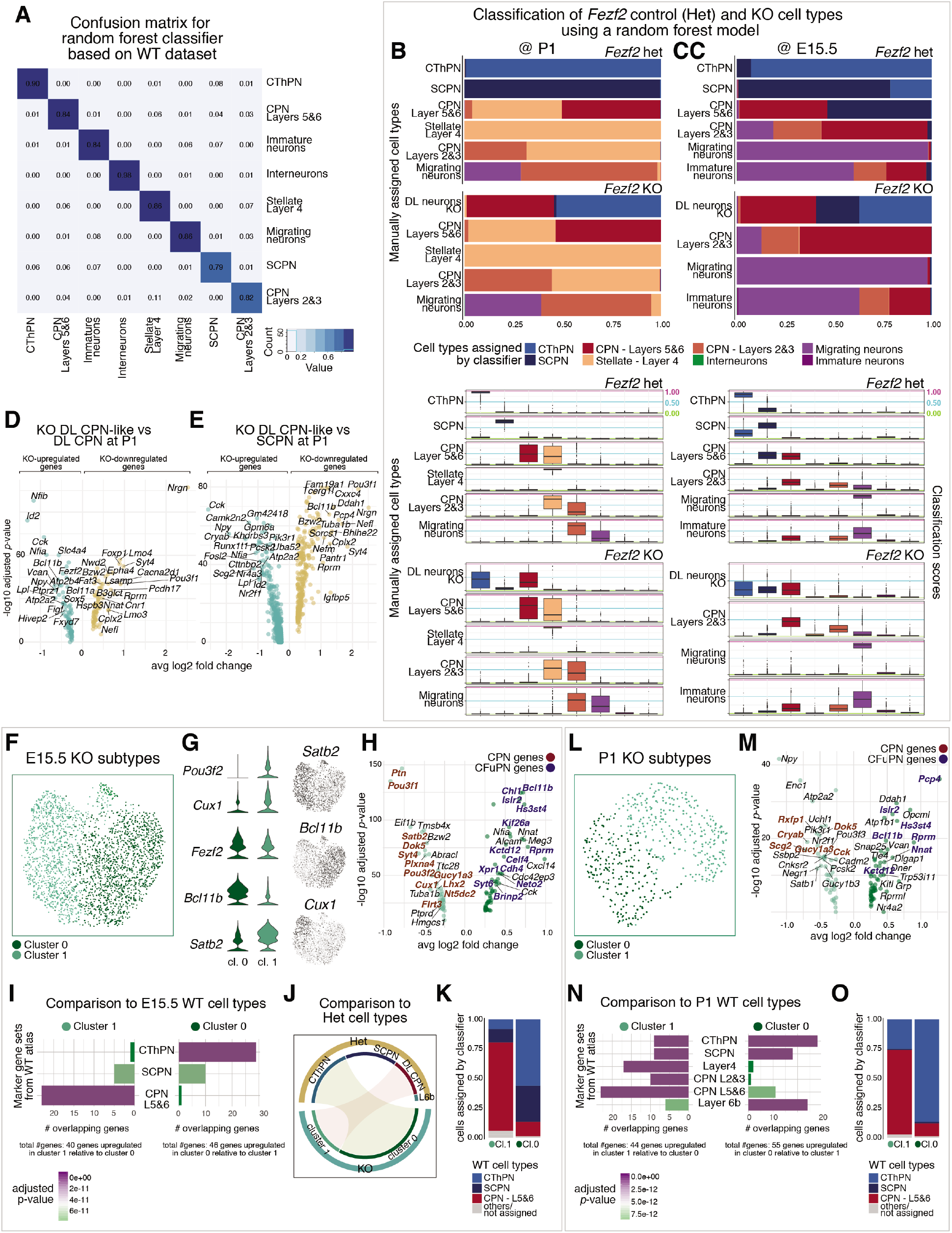
CFuPN acquire CThPN-like and Layers 5&6 CPN-like identities in the absence of *Fezf2*. **A** Confusion matrix for random forest classifier calculated using 1,000 cells per cluster of the WT developmental atlas. The remaining held-out cells were used to test accuracy. **B-C** Classification of control *(Fezf2* het) and *Fezf2* KO excitatory neurons by the classifier presented in **A**, for P1 (**B**) or E15.5 (**C**) datasets. Cells are grouped according to their manually assigned identity based on the expression of marker genes. Bottom panels show the corresponding classification scores. Lines in magenta, cyan and green indicate 1, 0.5 and 0 values, respectively. **D-E** Differential expression analysis of the aberrant layer 5&6 CPN-like cells from the KO-exclusive populations at P1 compared to layers 5&6 CPN (**D**) or SCPN (**E**) populations in the control. **F-H** Two subtypes of deep-layers KO cells were identified at E15.5. Subclustering of deep-layers KO-exclusive cells alone at E15.5 (**F**) shows a *Satb2^LOW^, Bcl11b^HIGH^* cluster (cluster 0), and a *Satb2^HIGH^* cluster expressing also CPN markers *Cux1* and *Pou3f2* (cluster 1), as indicated in the violin plots and UMAPs (**G**). Differential expression analysis between both subtypes indicates enrichment of CFuPN genes in cluster 0 and CPN genes in cluster 1 (**H**). **I** Comparison to neurons in E15.5 wild-type data showing overlap between differentially expressed genes and markers from E15.*5* neuronal subtypes (left). Bars indicate number of overlapping genes and are colored by the adjusted *p*-value calculated by hypergeometric test for significant enrichment. **J** Circular plot showing correlation between KO and control subpopulations of deep layers neurons. Cell types are indicated in the edge of the circle and shaded connected area indicates correlation of transcriptomes. **K** Classification of cells from both E15.5 KO-specific clusters according to random forest classifier shows good agreement between both annotations. **L-M** Sub-clustering (**L**) and differential expression analysis (**M**) of deep-layers KO-exclusive cells alone at P1 reveals two subpopulations that correspond to CThPN-like and layers 5&6 CPN-like populations. **N** Number of marker genes from KO clusters overlapping with markers from P1 wild-type neurons (similar to panel **I**). **O** Classification of cells from both P1 KO-specific clusters according to random forest classifier shows good agreement between both annotations.

**Supplementary information Figure 12 (related to Figure 4).**
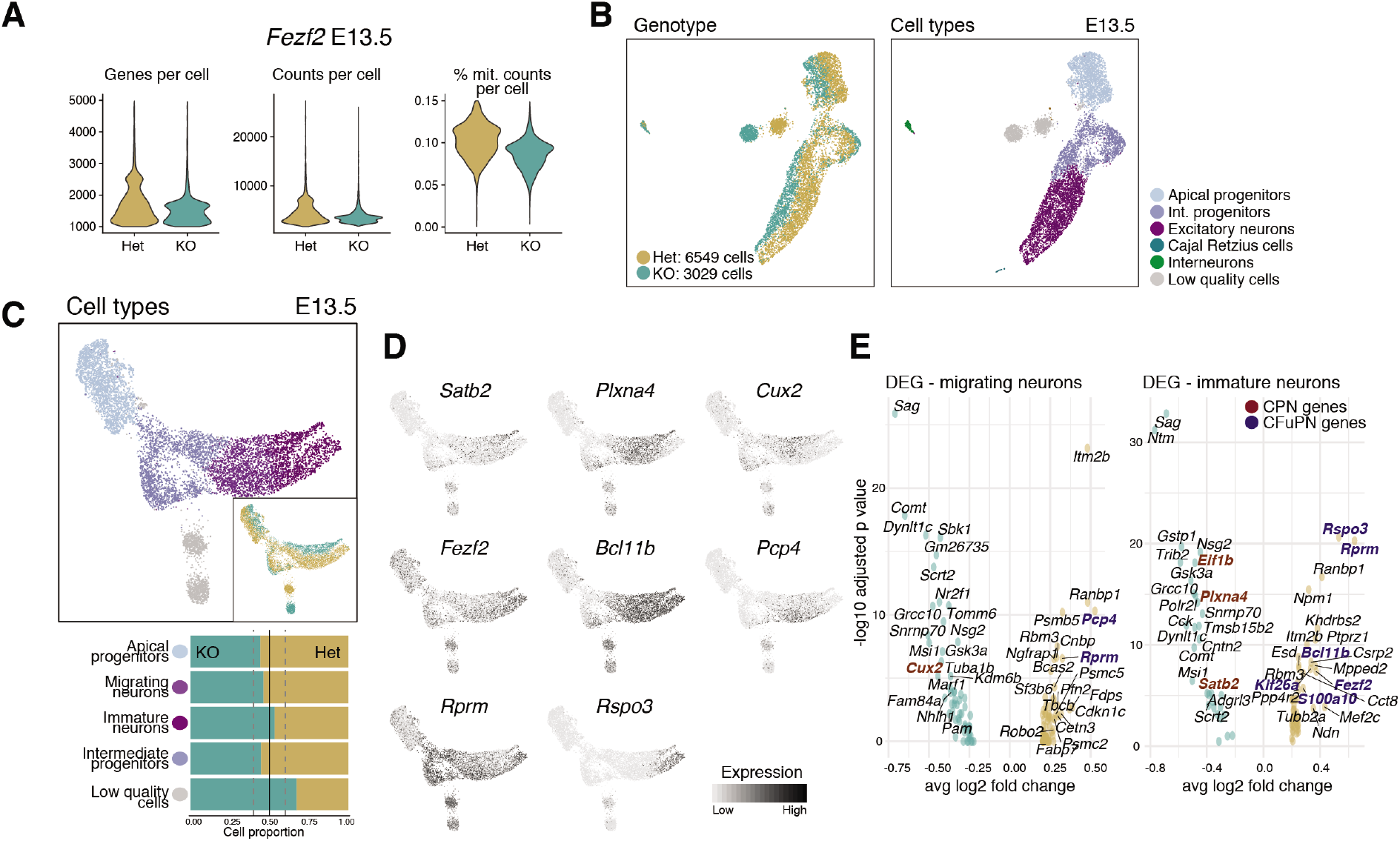
*Fezf2* acts post-mitotically to direct CFuPN differentiation. **A** Violin plots of number of genes (left) and mRNA molecules (counts; middle), and percentage of mitochondrial counts (right) per cell in control and KO *Fezf2* E13.5 single cell transcriptomes. **B** UMAP visualizations of combined control and KO complete data sets, colored by genotype or assigned cell type. **C** Dorsally-derived cells in *Fezf2* control and KO E13.5 scRNA-seq, visualized via UMAP and colored by cell types (top). Proportion of cells in each cell type, according to their genotype (bottom). **D** Expression of CPN *(Satb2, Plxna4, Cux2)* and CFuPN *(Fezf2, Bcl11b, Pcp4, Rprm, Rspo3)* marker genes overlaid on E13.5 *Fezf2* scRNA-seq data. **E** Differential expression analysis between control and KO migrating or immature neurons shows upregulation of a subset of CPN marker genes and downregulation of CFuPN-specific genes.

## Notes

### Competing Interest Statement

P.A. is a SAB member in System 1 Biosciences and Foresite Labs and is a co-founder of FL60. A.R. is a co-founder of and equity holder in Celsius Therapeutics, equity holder in Immunitas, and a SAB member of ThermoFisher Scientific, Syros Pharmaceuticlas, Asimov, and Neogene Therapeutics.

